# APOE isoforms shape brain network organization and its coupling with cardiac phenotypes in mice

**DOI:** 10.64898/2026.11.23.768680

**Authors:** Alexandra Badea, Kelly Shahu, Rohan Nadkarni, Jacques A Stout, Hae Sol Moon, Ali Mahzarnia, Zayar Han, Jessica Tremblay, Robert J Anderson, Matt Foster, Michael Lutz, Daniel Laskowitz, Cristian Badea

## Abstract

**Background:** Apolipoprotein E (APOE) isoforms influence Alzheimer’s disease susceptibility and cardiovascular physiology, but their roles in functional brain network organization and brain–heart relationships remain incompletely understood. We investigated whether APOE genotype modifies functional brain networks and their cardiovascular, behavioral, and molecular correlates.

**Methods:** Resting-state functional MRI was analyzed in 261 mice expressing human APOE2, APOE3, or APOE4, with 31 *Apoe*-knockout mice as additional comparators. Cardiac computed tomography, regional brain volume, intrinsic T1, cerebral blood flow (CBF), spatial-navigation behavior, and regional transcriptomics were evaluated in partially overlapping or independent cohorts. Covariate-adjusted analyses assessed network topology, regional connectivity, and APOE-dependent brain–cardiac associations, with false-discovery-rate correction within designated testing families.

**Results:** Compared with APOE2, APOE4 mice exhibited lower positive functional connectivity strength (β = −0.542, q = 0.0036), clustering (β = −0.488, q = 0.0041), and modularity (β = −0.375, q = 0.0252), together with higher global efficiency and altered persistent topology. Positive regional connectivity was lower in 15 cortical regions within the designated 244-region testing family, with left cingulate area 24b also surviving correction across all 1,464 regional tests (q = 0.0394). Among 108 mice with paired cardiac CT and functional MRI, APOE modified the association between left atrial volume and negative-network β₀ persistence (interaction p = 0.00386; q = 0.0463 across 12 persistence tests), predominantly reflecting a negative association in APOE3 mice; this interaction did not survive correction across all 28 global tests. Complementary regional MRI analyses identified genotype-dependent cardiac associations involving brain volume, intrinsic T1, and CBF, including three anatomically matched T1–CBF associations in limbic-related and pontine regions. APOE4 mice exhibited higher absolute winding number during spatial navigation and lower delayed spatial-memory probe occupancy. Cardiac ejection fraction was associated with acquisition trajectories in a smaller paired cohort. Independent cingulate transcriptomics identified APOE-associated lipid/cholesterol and inhibitory-neuron marker signatures (q < 0.05).

**Conclusions:** APOE isoforms are associated with distinct functional brain network architectures and genotype- dependent relationships between cerebral and cardiac phenotypes. Convergence across functional connectivity, regional MRI, behavior, and independent molecular measurements implicates distributed cortical and limbic- related systems, including the cingulate cortex. These findings identify candidate imaging phenotypes for investigating APOE-sensitive brain–heart relationships, without establishing their causal direction or underlying mechanisms.

## Introduction

The brain and heart interact through bidirectional neural, vascular, mechanical, and biochemical pathways that coordinate cardiovascular regulation and cerebral physiology (Valenza, Matić, and Catrambone 2025). Distributed cortical and subcortical networks, including the cingulate and insular cortices, hypothalamus, and brainstem, participate in autonomic cardiovascular control, while cardiac function influences cerebral perfusion and vascular physiology (Cechetto 2014). Large-scale human imaging studies have demonstrated associations between cardiac structure and function and brain morphology, microstructure, and functional connectivity. In more than 40,000 participants, Zhao et al. (Zhao et al. 2023) identified shared phenotypic and genetic associations between cardiac and brain MRI traits. These findings raise the question of whether genetic factors that influence neural organization also modify relationships between brain networks and cardiovascular phenotypes.

Apolipoprotein E (APOE) is a compelling candidate modifier because its major human isoforms, APOE2, APOE3, and APOE4, differentially influence lipid metabolism, neuroimmune responses, cerebrovascular integrity, and susceptibility to late-onset Alzheimer’s disease (Belloy, Napolioni, and Greicius 2019) (Safieh, Korczyn, and Michaelson 2019) (Yamazaki et al. 2019). APOE4-associated blood–brain barrier dysfunction suggests that genotype-dependent cerebrovascular processes may influence brain function independently of overt neurodegeneration (Montagne et al. 2020). Experimental studies using photon-counting cardiac CT in humanized APOE mice have also demonstrated genotype-dependent differences in cardiac anatomy and responses to exercise and dietary challenges (Allphin et al. 2023; Allphin et al. 2025). Together, these findings establish APOE as a modifier of both cerebral and cardiovascular phenotypes, but whether its isoforms alter relationships between functional brain organization and cardiac structure or function remains unclear.

Resting-state functional MRI provides a systems-level approach to examining these relationships by characterizing coordinated activity across distributed neural networks. Graph-theoretical measures quantify network connectivity, segregation, and integration, while persistent homology captures complementary topological properties across connectivity thresholds (Stolz, Harrington, and Porter 2017) (Songdechakraiwut and Chung 2020). Previous studies in humanized APOE mouse mode ls have identified genotype- and risk- factor-associated alterations in structural brain networks (Badea et al. 2019) (Winter et al. 2024) and resting- state functional connectivity (Zerbi et al. 2014). However, whether APOE-associated differences in functional network organization correspond to distinct cardiac phenotypes, and whether these relationships vary across APOE isoforms, remains largely unexplored. This question is particularly relevant to the central autonomic network, in which cortical regions interact with hypothalamic and brainstem structures involved in cardiovascular regulation (Cechetto 2014). Complementary measures of regional brain volume, intrinsic T1 relaxation, and cerebral blood flow may provide additional anatomical, tissue-sensitive, and vascular context for functional network differences. Behavioral and molecular phenotypes can further inform their biological relevance. Nevertheless, associations between separately acquired brain and cardiac measurements cannot distinguish neural regulation of the heart from cardiovascular influences on the brain or shared systemic mechanisms.

Here, we investigated how human APOE isoforms shape functional brain network organization and its cardiovascular, behavioral, and molecular correlates in mice. We first characterized global network topology and regional functional connectivity across humanized APOE2, APOE3, and APOE4 mice, with *Apoe*-knockout mice as complementary comparators. We then tested whether associations between cardiac CT phenotypes and functional network measures differed by APOE genotype in animals with paired measurements. In complementary cohorts, we examined genotype-dependent associations between regional brain MRI biomarkers and cardiac phenotypes, spatial learning and memory, and independent regional transcriptomic and proteomic signatures. We hypothesized that APOE isoforms would be associated with distinct functional network architectures and genotype-dependent brain–cardiac relationships. By integrating these complementary measurements, we sought to identify anatomically convergent patterns and candidate biological substrates relevant to APOE-sensitive brain–heart interactions.

## Methods

### Animals and experimental design

Mice homozygous for human APOE2, APOE3, or APOE4 were studied on either the endogenous mouse *Nos2* background or a humanized NOS2 background (HN; HuNOS2tg/mNos2−/−). *Apoe*-knockout mice on the mouse *Nos2* background served as additional comparators. Animals received standard chow or a high-fat diet containing 45% kcal from fat (Research Diets, D12451i) for approximately four months before MRI. Selected cohorts also had access to running wheels. Age at MRI was calculated from verified birth and acquisition dates. All procedures were approved by the Duke University Institutional Animal Care and Use Committee.

The study included 261 humanized APOE and 31 *Apoe*-knockout mice. Cardiac imaging, behavioral assessment, regional quantitative MRI, and molecular profiling were performed in partially overlapping or independent cohorts, with eligibility determined separately for each analysis (Figure 1; Supplementary Methods). Animal identity, imaging dates, genotype, experimental exposures, and relevant covariates were reconciled before multimodal analyses. The overall study flow is described in Figure 1

**Figure 1.**
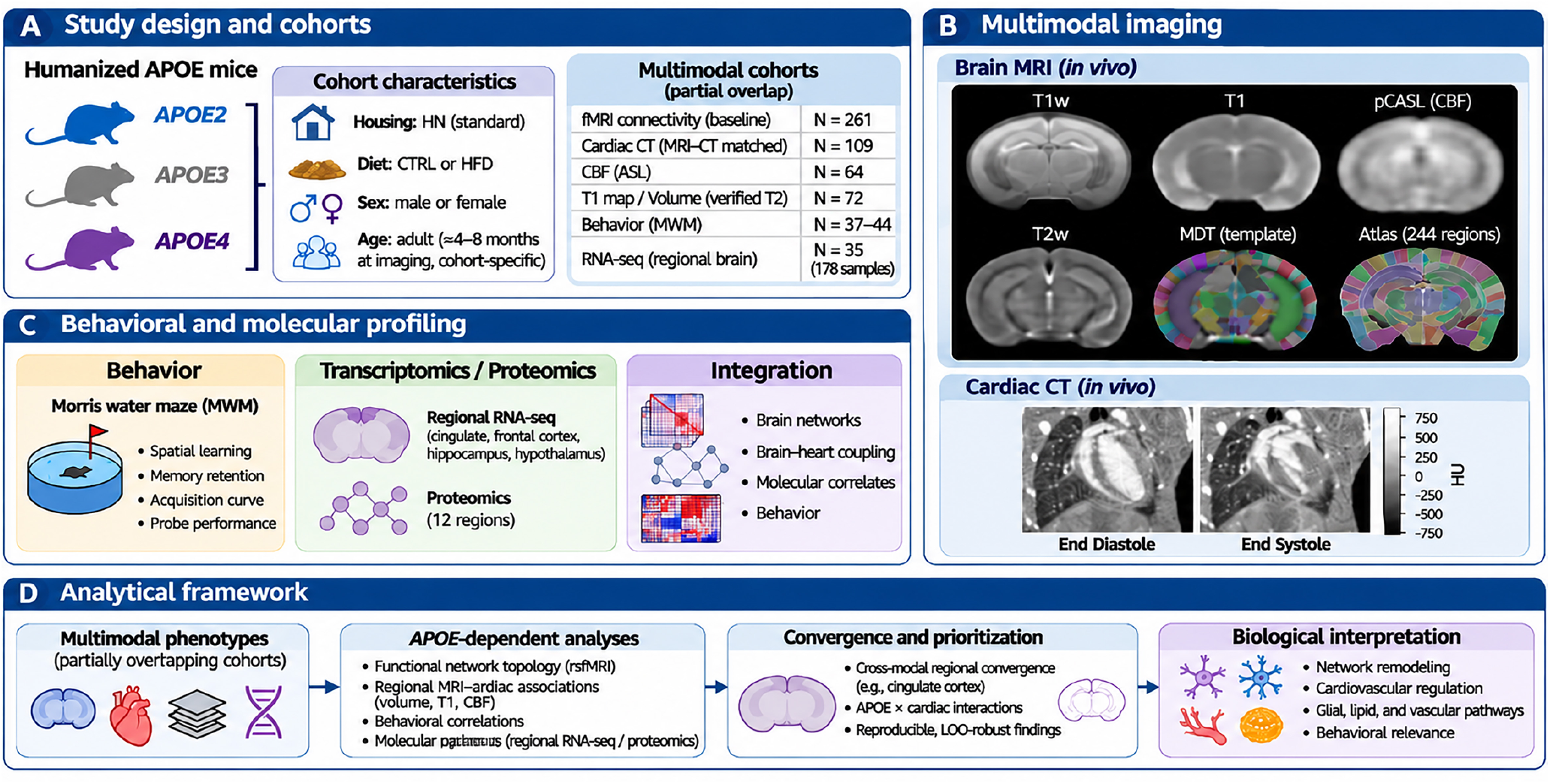
Study design, multimodal characterization, and analytical framework in humanized APOE mice. (A) Study design and cohorts. Humanized APOE2, APOE3, and APOE4 mice were characterized across partially overlapping cohorts differing in housing conditions, diet, sex, and age. The functional connectome dataset comprised 292 mice (261 humanized APOE mice and 31 *Apoe*-knockout mice). Additional cohorts included 108 mice with paired resting-state functional MRI (rsfMRI) and cardiac CT, 130 mice with Morris water maze (MWM) measurements, and 35 mice contributing 178 regional RNA-seq samples. Regional MRI–cardiac analyses included 92 mice for regional volume (APOE2, n = 36; APOE3, n = 30; APOE4, n = 26) and 64 mice each for intrinsic T1 and cerebral blood flow (CBF; APOE2, n = 35; APOE3, n = 21; APOE4, n = 8). These modality- specific cohorts partially overlapped and were distinct from the paired cardiac–functional connectome cohort. (B) Multimodal imaging. Representative coronal brain images illustrate T1-weighted (T1w), quantitative T1, pseudo-continuous arterial spin labeling (pCASL)-derived CBF, and T2-weighted (T2w) MRI, together with the minimum deformation template (MDT) and anatomical atlas used for regional analyses. Contrast-enhanced cardiac CT images at end diastole and end systole illustrate measurements of cardiac chamber volumes and function, including stroke volume, ejection fraction, and cardiac output. (C) Behavioral and molecular profiling. Spatial learning and memory were assessed using the MWM. The nested cardiac–behavior analysis included 37 mice in the primary cohort and 44 in an overlapping sensitivity cohort. Regional RNA-seq comprised 178 samples from 35 mice, including 32 mice contributing cingulate cortex samples. Transcriptomic and exploratory proteomic measurements provided complementary molecular context involving lipid metabolism, glial function, inhibitory signaling, and vascular regulation. (D) Analytical framework. Multimodal analyses characterized APOE-dependent functional network organization, genotype-modified associations between regional brain MRI biomarkers (volume, T1, and CBF) and cardiac phenotypes, and relationships with behavioral performance. Cross-modal anatomical convergence, genotype interactions, and sensitivity analyses were used to identify candidate regions and biological systems, including the cingulate cortex. Molecular signatures from independent cohorts informed biological interpretation without establishing molecular mediation or causal brain–heart relationships. Cohort sizes reflect modality-specific availability, eligibility, and quality-control criteria.

### MRI acquisition and preprocessing

Brain MRI was performed at 7 T using a Bruker BioSpec 70/20 scanner (ParaVision 6.0.1) equipped with a 1,000 mT/m gradient insert, volume transmit coil, and four-channel receive array. Animals were anesthetized with approximately 1.5% isoflurane in air, with physiological monitoring and circulating-water warming. Structural MRI included three-dimensional T1-weighted images at 0.1-mm isotropic resolution or multislice T2-weighted images at 0.156 × 0.156 × 0.55 mm resolution, depending on acquisition cohort.

Resting-state functional MRI used coronal echo-planar imaging with a 64 × 50 acquisition matrix, 19.2 × 15 mm field of view, 0.3-mm slice thickness, and repetition time of approximately 2.25 s for historical multiecho acquisitions. Echo times varied across acquisition protocols; representative values were 5.0, 19.3, and 33.6 ms. Single-echo acquisitions, where applicable, were identified separately in the acquisition metadata. Acquisition protocol and scan year were included as covariates in functional-connectivity analyses where estimable.

Brain masks were generated using RS²-Net (Lin et al. 2024) and reviewed for anatomical accuracy. Anatomical image processing and atlas-based regional analysis followed principles established in the Small Animal Multivariate Brain Analysis framework (SAMBA; (Anderson et al. 2019)). A minimum-deformation template derived from manganese-enhanced T1-weighted images and incorporating the symmetric CHASS mouse brain atlas was registered to individual anatomical images using Advanced Normalization Tools (ANTs) (Avants et al. 2011), with rigid, affine, and symmetric normalization transformations.

Functional images were processed in native echo-planar imaging space. Preprocessing included slice-timing correction, motion correction, censoring of frames with framewise displacement greater than 0.1 mm, multiecho combination where applicable, and boundary-based registration to individual anatomy. Nuisance regression included motion parameters and their derivatives, five anatomical CompCor components each from white matter and cerebrospinal fluid {Behzadi, 2007 #21), and polynomial trends, followed by temporal filtering between 0.01 and 0.1 Hz. Regional time series were extracted using fractional atlas-label weights over a common 450-s analysis window. Detailed acquisition parameters and quality-control procedures are provided in the Supplementary Methods.

Manganese-enhanced anatomical imaging and the intrinsic quantitative T1 measurements used in the regional MRI analyses were derived from distinct acquisition protocols and were analyzed separately.

Cerebral blood flow (CBF) was measured using pseudo-continuous arterial spin labeling (pCASL), following previously established methods for high-field rodent perfusion imaging {Hirschler, 2018 #27} (Hirschler et al. 2018) and our earlier application in the CVN-AD mouse model (Badea et al. 2025). Acquisition parameters were identical to those previously reported: 30 control–label pairs, repetition time/echo time (TR/TE) = 4500/16.43 ms, labeling duration = 3000 ms, and post-labeling delay = 300 ms. Quantitative CBF maps were registered to individual anatomical images and regional perfusion values extracted using the CHASS mouse brain atlas. Regional CBF measurements were analyzed in 64 humanized APOE mice (APOE2, n = 35; APOE3, n = 21; APOE4, n = 8) with matched cardiac CT measurements. These measurements were acquired without manganese enhancement and analyzed separately from intrinsic T1 relaxation and regional brain volume.

### Functional connectivity and network analysis Connectome construction and anatomical coverage

Functional-connectivity analyses included 292 mice (APOE2, n = 109; APOE3, n = 69; APOE4, n = 83; *Apoe*- knockout, n = 31). Primary comparisons among humanized APOE genotypes were restricted to 261 animals.

Regional functional connectivity was calculated from the extracted time series and represented as Fisher-z- transformed correlation matrices. Of the 324 regions in the symmetric CHASS atlas, 260 gray-matter regions were initially selected after excluding white matter, cerebrospinal fluid, and ventricular compartments. An anatomical coverage criterion, independent of genotype and connectivity outcome, retained regions represented by at least one EPI-voxel equivalent in at least 95% of quality-controlled mice. The resulting 244-region atlas yielded 29,646 unique undirected connections per animal. Primary connectome models adjusted for age at MRI, sex, diet, MRI-associated body mass, HN status, acquisition protocol, and scan year, where estimable. Exercise assignment was excluded because its recorded timing did not establish exposure before imaging.

### Global graph measures and statistical inference

Nine global network outcomes were evaluated: mean positive and negative connectivity strength, global efficiency, clustering coefficient, modularity, and zero- and one-dimensional persistent-homology area-under- the-curve measures for positive and negative connectivity. Network measures were rank-transformed and evaluated using covariate-adjusted regression. Standardized coefficients describe effects on the transformed outcome scale and should not be interpreted as absolute or percentage changes in connectivity.

Primary genotype comparisons included APOE4 versus APOE2, APOE3 versus APOE2, and APOE4 versus APOE3. Covariate-adjusted permutation inference used a Freedman–Lane-style procedure with 5,000 permutations, and uncertainty was estimated using 2,000 bootstrap replicates. Benjamini–Hochberg false- discovery-rate (FDR) correction was applied within the designated testing families. Knockout-inclusive comparisons, risk-factor analyses, and ordered genotype contrasts were evaluated in their respective testing families. Because *Apoe*-knockout animals were HN-negative and acquired within a restricted protocol and scan- year combination, knockout comparisons were interpreted with these acquisition constraints in mind.

### Multivariate connectome analysis

Whole-connectome genotype separation was evaluated using cross-validated distances between covariate- adjusted connectivity-effect patterns. For each of 200 random splits, genotype groups were divided into independent halves, and cross-validated squared distances were estimated from cross-products of independently derived genotype contrasts across the 29,646 edges. Distances were averaged across splits.

Classical multidimensional scaling was applied to the nonnegative distance representation. Nonpositive cross- validated squared-distance estimates, which may arise from sampling variability, were treated as indicating no detectable separation for visualization. Uncertainty was characterized using 500 within-group bootstrap embeddings aligned by Procrustes transformation. A permutation test with 999 genotype-label permutations evaluated overall multivariate separation among the humanized APOE mice.

Principal component analysis was also performed on nine global network measures after removing covariate- associated variation while retaining estimated genotype-related effects. Standardized residualized measures were decomposed using singular value decomposition. Genotype centroids and bootstrap uncertainty ellipses were displayed in the first two principal components.

### Persistent homology

Persistent homology was used to characterize network topology across proportional connectivity thresholds (Stolz, Harrington, and Porter 2017) (Songdechakraiwut and Chung 2020). Positive and negative edges were analyzed separately, with negative edges ranked by absolute correlation magnitude. Clique-complex filtrations were constructed over increasing edge density, and persistence diagrams were computed using Ripser (Bauer 2021).

Zero-dimensional features (β₀) represented connected components, whereas one-dimensional features (β₁) represented topological cycles. Persistence area under the curve (AUC) was calculated from feature lifetimes across the full filtration, truncated at edge density 1.0. Betti curves were evaluated using a fine density grid with increments of 0.0005 through density 0.10 and a coarser grid thereafter. Covariate-adjusted genotype curves were accompanied by pointwise 95% bootstrap intervals derived from 2,000 within-genotype resamples. Statistical comparisons used full-filtration persistence AUC rather than pointwise tests at individual connectivity thresholds.

### Regional connectivity analysis

Regional positive and negative nodal strength were calculated separately from the symmetric, diagonal-zeroed Fisher-z connectivity matrices. For each of the 244 regions, positive strength was defined as the mean positive Fisher-z connection magnitude across the remaining 243 nodes; negative strength was defined analogously using absolute negative Fisher-z magnitudes.

Within the 261 humanized APOE mice, regional strength was rank-transformed and modeled using covariate- adjusted ordinary least squares. The three pairwise APOE contrasts were evaluated separately for positive and negative connectivity, yielding six predefined families of 244 tests (1,464 tests in total). Two-sided regression t- test P values were adjusted using Benjamini–Hochberg correction within each contrast-by-connectivity-sign family. As a more conservative sensitivity analysis, FDR correction was also applied across all 1,464 regional tests.

### Cardiac CT and brain–heart network associations Cardiac imaging and cohort linkage

Cardiac structure and function were quantified using contrast-enhanced photon-counting micro-computed tomography (PCCT), following previously established methods for cardiac phenotyping in humanized APOE mouse models (Allphin et al. 2023) (Allphin et al. 2025). Cardiac measurements included chamber volumes, myocardial volumes, ventricular ejection fraction, stroke volume, and cardiac output.

Cardiac CT measurements were linked to resting-state functional-connectivity data using a curated animal-level identity crosswalk. The primary paired cohort comprised 108 control-diet humanized APOE mice (APOE2, n = 46; APOE3, n = 27; APOE4, n = 35), with cardiac CT and brain MRI acquired no more than 90 days apart. This cohort included 80 historical and 28 newer acquisitions, analyzed together. *Apoe*-knockout mice were excluded from these analyses.

Primary cardiac predictors were diastolic left atrial volume, diastolic myocardial volume, left-ventricular ejection fraction, and cardiac output. Historical diastolic left atrial measurements were harmonized with maximum left atrial volume in the newer CT dataset. Volumes recorded in microliters were converted to milliliters when necessary. Cardiac volumes were analyzed without normalization to body mass, which was included as a model covariate.

### Global network–cardiac interactions

The principal cardiac–network analysis evaluated the negative-connectivity β₀ persistence AUC. For each cardiac phenotype, the model included APOE genotype, the cardiac measure, and their interaction, adjusting for sex, age at MRI, signed CT–MRI interval, CT-day body mass, HN status, acquisition cohort, scan year, and imaging protocol.

Cardiac predictors were standardized using the eligible 138-animal cardiac–connectome cohort, and persistence outcomes were rank-based inverse-normal transformed using the same reference cohort. Primary models were subsequently fitted in the 108 control-diet animals.

Ordinary least squares regression with HC3 heteroskedasticity-robust covariance was used for estimation. APOE-dependent cardiac associations were evaluated using joint two-degree-of-freedom Wald tests. Genotype- specific slopes, pairwise slope contrasts, and corresponding 95% confidence intervals were obtained as linear combinations of fitted regression coefficients.

Benjamini–Hochberg correction was applied across four cardiac predictors for the primary topology outcome, across 12 persistence-related cardiac tests, and across the complete family of 28 global cardiac–network tests. These nested families distinguished focused from broader multiplicity control. Because the four cardiac predictors were selected following exploratory screening, the broader correction was also considered when interpreting the findings.

Sensitivity analyses excluded body mass from the regression model and repeated the primary analysis after sequentially removing each animal.

### Regional network–cardiac interactions

APOE-dependent cardiac associations were additionally evaluated for positive and absolute negative nodal connectivity strength in each of the 244 regions. For each cardiac phenotype, 488 regional outcomes were examined.

The four primary cardiac phenotypes yielded 1,952 regional interaction tests. An expanded exploratory analysis included 14 cardiac phenotypes, resulting in 6,832 tests. Benjamini–Hochberg correction was applied within each 488-test cardiac family, across the original 1,952 tests, and across the expanded 6,832-test family.

Additional exploratory analyses restricted the regional search to the 20, 40, or 80 largest atlas regions, selected using independent reference-anatomical volume rankings. Selected findings were evaluated in the historical 80- mouse subset and by leave-one-animal-out analysis. The historical subset was treated as a nested sensitivity cohort rather than an independent replication sample.

### Spatial navigation and brain–heart–behavior associations Behavioral assessment and absolute winding number

Spatial learning and memory were evaluated using the Morris water maze in 130 humanized APOE mice (APOE2, n = 47; APOE3, n = 44; APOE4, n = 39), following established spatial-learning procedures (Vorhees and Williams 2006). Acquisition testing comprised four trials per day across five days, yielding 2,600 trial observations.

Spatial navigation trajectories were characterized using absolute winding number (AWN), a path-based measure of navigational complexity and search strategy previously developed and evaluated in humanized APOE mice (Badea et al. 2022). Unlike conventional escape latency or path length, AWN quantifies the cumulative winding of the swimming trajectory and provides complementary information about the spatial organization of search behavior. AWN was analyzed following log(1 + AWN) transformation. Total swimming distance was analyzed separately using the same transformation. Spatial-memory retention was assessed using target-quadrant occupancy during Day 5 and Day 8 probe tests. Behavioral models included APOE genotype and relevant biological covariates, including HN status, sex, diet, and age. Genotype differences in acquisition were evaluated using overall APOE effects and genotype-by-day interactions, with acquisition day modeled as either a continuous or categorical variable. Paired probe models evaluated genotype, probe day, and their interaction, accounting for repeated measurements within mice. Pairwise genotype comparisons were corrected for multiple testing within the designated families.

### Functional connectivity and spatial navigation

Regional positive and negative nodal-strength measures were matched to behavioral observations using verified animal identifiers. An unbiased regional screen examined 244 regions, two connectivity signs, and three acquisition endpoints (Day 1, Day 5, and learning rate), yielding 1,464 tests. Benjamini–Hochberg correction was applied across the full screening family. Targeted longitudinal analyses subsequently evaluated negative connectivity in left hypothalamic PVG and left cingulate area 30, and positive connectivity in left cingulate area 25. Regional connectivity measures were standardized before modeling. Connectivity-by-day associations were estimated using repeated-trial models with mouse-clustered uncertainty, adjusting for genotype and relevant biological covariates. Multiple-testing correction for these targeted analyses was maintained separately from the unbiased regional screen.

### Cardiac function and spatial navigation

Cardiac ejection fraction was linked to behavioral records using an audited CT–behavior identity crosswalk. The primary conservative cohort comprised 37 animals. A sensitivity cohort of 44 animals additionally included seven adjudicated identity matches; these overlapping cohorts were not considered independent replications.

Ejection fraction was standardized within the analysis cohort. Longitudinal models evaluated associations between ejection fraction and acquisition-day changes in log-transformed AWN while adjusting for APOE genotype and biological covariates. Mouse-clustered uncertainty estimates accounted for repeated behavioral trials. An additional exploratory model examined the joint association of APOE genotype, left cingulate area 25 positive connectivity, and ejection fraction with navigation performance. The model included all lower-order and three-way interaction terms, HN status, sex, diet, standardized age, centered acquisition day, and trial number. Inference was evaluated at trial, mouse-day, and mouse levels. A null-restricted Rademacher wild-cluster bootstrap with 399 replicates was used to assess the joint three-way interaction. Six targeted genotype-specific contrasts were adjusted using the Holm procedure. Exploratory analyses of acquisition target-quadrant path fraction were derived from individual tracking records, with unavailable or invalid trajectories treated as missing.

### Regional molecular mapping and transcriptomic analysis Imaging-derived regional connectivity maps

Regional APOE4-versus-APOE2 differences in positive nodal strength were estimated from the 244-region functional-connectivity atlas using the primary imaging covariate model. These imaging-effect maps were compared with regional gene-expression data from the Allen Mouse Brain Atlas, comprising 4,082 assayed genes (Wang et al. 2020) (Lein et al. 2007). Anatomical correspondence was evaluated using both manually curated CHASS-to-Allen mappings and registration-based mapping to the Allen Common Coordinate Framework. The primary regional comparison included 89 bilateral-averaged anatomical structures.

### Spatial gene-expression associations

Gene-expression analyses evaluated predefined neuronal and glial cell-type markers, Gene Ontology pathways, and Alzheimer’s disease risk-gene orthologs. For each gene set, regional gene-wise Spearman correlations were calculated between reference-atlas expression and the APOE4-versus-APOE2 nodal-strength effect map. Inference incorporated spatially autocorrelated surrogate imaging maps and size-matched random-gene-set null distributions, with 5,000 surrogate iterations of each type. Combined significance required nominal support from both spatial and gene-set null tests. Benjamini–Hochberg correction was applied within the designated mapping and gene-set testing families. Regional composite expression scores used for visualization were distinguished from the gene-wise correlations used for statistical inference.

### Regional RNA sequencing and proteomics

Laboratory RNA sequencing included 178 samples from 35 humanized APOE mice across 12 brain regions. Expression profiles were anatomically matched to corresponding Allen reference regions and to an exploratory proteomics dataset covering 12 regions from two APOE4/HN mice. For cross-platform comparisons, shared gene-expression and protein-abundance profiles were standardized within the corresponding data source. Regional expression concordance was evaluated using Spearman correlations for selected excitatory-neuron markers (*Slc17a7, Camk2a, Tbr1, Nrgn,* and *Slc17a6*) and a synaptic Gene Ontology signature. These comparisons characterized regional molecular organization rather than replication of genotype effects.

### Cingulate transcriptomic signatures

Genotype-dependent molecular signatures were evaluated in cingulate cortex RNA-seq samples from 32 mice (APOE2, n = 11; APOE3, n = 14; APOE4, n = 7). Seventeen predefined cell-type marker and biological-pathway signatures were analyzed using regression models including categorical APOE genotype, HN status, sequencing or processing order, sex, and diet. Overall genotype effects were evaluated using two-degree-of-freedom omnibus tests. Pairwise genotype contrasts were estimated with HC3 heteroskedasticity-robust covariance and 95% confidence intervals. Benjamini–Hochberg correction was applied separately across the 17 omnibus tests and 51 pairwise genotype contrasts. Sensitivity analyses were performed in the 29 HN-negative animals and through leave-one-animal-out refitting. Because the RNA-seq and functional imaging measurements were derived from different animals, molecular signatures were interpreted as independent biological context rather than evidence of within-animal molecular mediation.

### Regional brain MRI biomarkers and cardiac associations Cohorts and quantitative MRI measures

Regional brain volume, intrinsic T1 relaxation, and cerebral blood flow (CBF) were evaluated in partially overlapping MRI–cardiac cohorts. Regional volume analysis included 92 animals (APOE2, n = 36; APOE3, n = 30; APOE4, n = 26). Intrinsic T1 and CBF analyses each included 64 animals (APOE2, n = 35; APOE3, n = 21; APOE4, n = 8). MRI and CT measurements were linked using verified animal identifiers and acquisition records. Regional volumes were normalized to total brain volume excluding exterior compartments, whereas intrinsic T1 and CBF were retained as quantitative regional measures. Intrinsic T1 was derived from acquisitions without manganese enhancement and was analyzed separately from manganese-enhanced structural imaging.Region definitions were based on verified CHASS atlas labels. Full-atlas analyses and analyses restricted to 244 connectome-compatible regions were treated as overlapping sensitivity analyses with distinct multiple-testing families, not independent replication datasets.

### APOE-dependent regional MRI–cardiac associations

For each regional MRI measure and cardiac phenotype, regression models included APOE genotype, the cardiac predictor, and an APOE-by-cardiac interaction. The primary HN-harmonized models included age, sex, exercise assignment, and humanized NOS2 status, with diet included when variable within the complete-case cohort. Body mass was not included in the primary models. A separate sensitivity analysis replaced exercise assignment with CT-associated body mass while retaining HN adjustment (M4 specification). APOE-dependent associations were evaluated using joint two-degree-of-freedom Wald tests with HC3 heteroskedasticity-robust covariance. Genotype-specific cardiac slopes and corresponding 95% confidence intervals were derived from the fitted models. Benjamini–Hochberg correction was applied separately within each MRI modality’s full region- by-cardiac interaction family. A corresponding analysis restricted to the 244 connectome-compatible regions provided an atlas-aligned sensitivity assessment. Regional APOE main effects were evaluated separately from genotype-by-cardiac interactions to distinguish differences in average MRI measurements from genotype- dependent brain–cardiac covariation.

### Cross-modality anatomical convergence

Multimodal convergence was evaluated by matching anatomical atlas regions and cardiac phenotypes across volume, T1, and CBF analyses. A regional MRI–cardiac pair was classified as overlapping when the APOE-by- cardiac interaction met the modality-specific FDR threshold (q < 0.05) in at least two modalities. Tested nonsignificant associations were distinguished from pairs unavailable for evaluation. Because modality cohorts were partially overlapping, cross-modality agreement was considered anatomical convergence rather than independent statistical replication. Associations were interpreted without assigning causal direction or inferring a common molecular mechanism.

### General statistical considerations

Analyses were conducted at the animal level or incorporated mouse-clustered inference for repeated behavioral observations. Covariate adjustment, outcome transformations, and multiple-testing families were specified for each analysis, reflecting differences in experimental design and hypothesis structure.

Unless otherwise stated, significance was evaluated using two-sided tests and Benjamini–Hochberg-adjusted q < 0.05. Holm correction was applied to designated pairwise behavioral contrasts. Interaction tests were distinguished from genotype-specific slopes, and nominal significance was not interpreted as family-wise corrected evidence.

Cohorts sharing animals, anatomical regions, or measurement sources were not treated as independent replications. Imaging, cardiac, behavioral, and molecular associations were interpreted as observational and did not establish causal direction, autonomic mediation, or direct molecular mechanisms.

## Results

### APOE-associated global network organization and topology APOE4 is associated with altered positive-network organization

In the 261 humanized APOE mice, APOE4 relative to APOE2 was associated with lower positive mean connectivity strength (β = −0.542, p = 0.0004, q = 0.0036), clustering (β = −0.488, p = 0.0006, q = 0.00405) and modularity (β = −0.375, p = 0.0056, q = 0.0252). Global efficiency was higher (β = +0.540, p = 0.0004, q = 0.0036). These results were significant within the archived main genotype-model FDR family (Table 1). The effects are coherent with an altered balance between local connectivity/segregation and graph-wide integration, but the direction of an individual network measure does not by itself establish improved or impaired neural function.

**Table 1.** Covariate-adjusted global positive-network genotype contrasts (source: Figure 2 tables document, Table 2)

| **Contrast** | **Positive strength** | **Clustering** | **Modularity** | **Global efficiency** | **Positive β<sub>1</sub> AUC** | **Positive β<sub>0</sub> AUC** |
|---|---|---|---|---|---|---|
| **APOE4 vs APOE2** | -0.54 (-0.84, -0.24); p 0.0004; q 0.0036 | -0.49 (-0.75, -0.22); p 0.0006; q 0.00405 | -0.38 (-0.65, -0.10); p 0.0056; q 0.0252 | +0.54 (+0.26, +0.81); p 0.0004; q 0.0036 | +0.42 (+0.15, +0.67); p 0.0024; q 0.013 | -0.54 (-0.81, -0.26); p 0.0004; q 0.0036 |
| **APOE3 vs APOE2** | -0.30 (-0.59, +0.00); p 0.0602; q 0.135 | -0.25 (-0.55, +0.04); p 0.0908; q 0.185 | -0.34 (-0.62, -0.05); p 0.0226; q 0.0678 | +0.24 (-0.07, +0.55); p 0.123; q 0.185 | +0.23 (-0.03, +0.51); p 0.102; q 0.185 | -0.35 (-0.65, -0.06); p 0.0222; q 0.0678 |
| **APOE4 vs APOE3** | -0.24 (-0.57, +0.06); p 0.124; q 0.185 | -0.24 (-0.53, +0.05); p 0.108; q 0.185 | -0.04 (-0.34, +0.25); p 0.793; q 0.807 | +0.30 (+0.01, +0.59); p 0.0502; q 0.123 | +0.19 (-0.09, +0.47); p 0.194; q 0.262 | -0.19 (-0.49, +0.10); p 0.236; q 0.303 |
| **KO vs APOE2** | -0.76 (-1.33, -0.16); p 0.0036; q 0.0324 | -0.63 (-1.20, -0.02); p 0.0084; q 0.0378 | -0.44 (-1.00, +0.13); p 0.0646; q 0.0969 | +0.59 (+0.01, +1.17); p 0.0168; q 0.0441 | +0.54 (-0.07, +1.11); p 0.0196; q 0.0441 | -0.49 (-1.03, +0.04); p 0.0506; q 0.0911 |
| **KO vs APOE4** | -0.23 (-0.78, +0.33); p 0.359; q 0.952 | -0.15 (-0.69, +0.43); p 0.506; q 0.952 | -0.07 (-0.60, +0.49); p 0.761; q 0.952 | +0.06 (-0.50, +0.59); p 0.787; q 0.952 | +0.14 (-0.44, +0.65); p 0.527; q 0.952 | +0.04 (-0.43, +0.57); p 0.854; q 0.952 |
| **ordered trend** | -0.26 (-0.40, -0.12); p 0.0002; q 0.0006 | -0.23 (-0.36, -0.09); p 0.0002; q 0.0006 | -0.17 (-0.30, -0.04); p 0.007; q 0.0105 | +0.24 (+0.10, +0.37); p 0.0002; q 0.0006 | +0.19 (+0.06, +0.32); p 0.0018; q 0.00324 | -0.23 (-0.36, -0.09); p 0.0006; q 0.00135 |

The APOE3-versus-APOE2 differences were generally intermediate in the same direction, but none of the six positive-network outcomes in Table 1 met q < 0.05 for this comparison; modularity (β = −0.34, q = 0.0678) and positive β₀ AUC (β = −0.35, q = 0.0678) were suggestive only. No displayed positive-network measure passed FDR correction for APOE4 versus APOE3. A prespecified ordered group contrast showed a consistent directional pattern across the six displayed network measures (all reported q ≤ 0.0105), but it does not imply a linear biological dose response and requires caution because the KO group has limited acquisition overlap.

### Persistent topology corroborates the positive-network differences

For positive connectivity, APOE4 compared with APOE2 showed lower β₀ persistence AUC (β = −0.537, p = 0.0004, q = 0.0036) and higher β₁ persistence AUC (β = +0.416, p = 0.0024, q = 0.0130). The Betti curves (Figure 2D–G) display how these differences evolve over proportional edge density; their insets summarize full- filtration AUC tests. The corresponding negative-connectivity persistence-AUC genotype contrasts did not reach FDR significance in the finalized analysis (Supplementary Table S2). These findings reflect altered graph topology across a rank-based filtration; β₁ cycles should not be equated with specific anatomical or physiological feedback loops.

**Figure 2.**
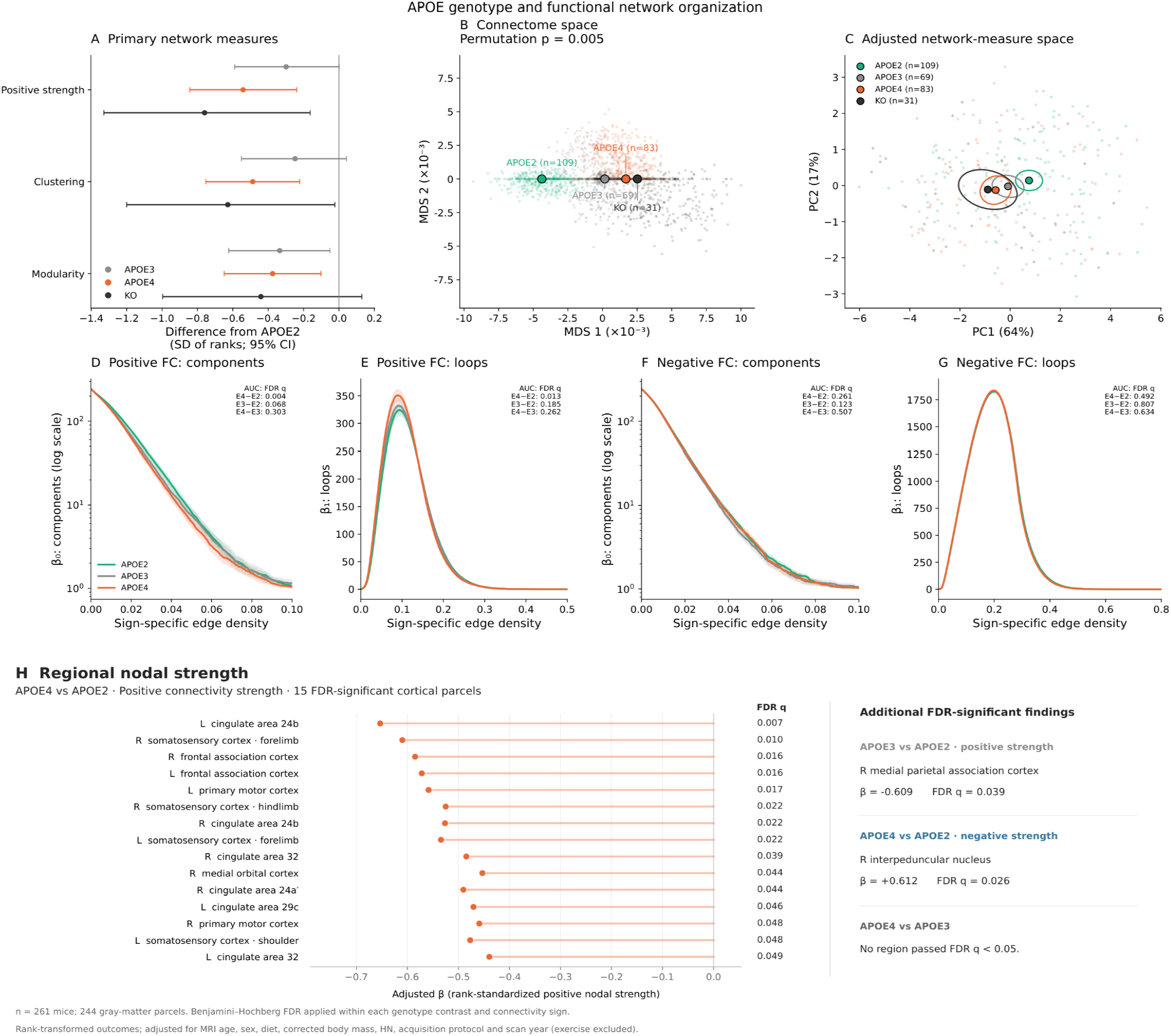
APOE genotype and functional network organization. (A) Adjusted contrasts of positive strength, clustering and modularity relative to APOE2 (95% confidence intervals). (B) Cross-validated whole-connectome MDS; the humanized genotype permutation p=0.005. (C) Adjusted nine-measure PCA, with centroids and uncertainty ellipses. (D–G) Positive and negative β₀ and β₁ Betti curves over proportional edge density; annotations reflect full-filtration AUC tests. (H) Covariate-adjusted APOE4–APOE2 positive-nodal-strength coefficients for the 15 cortical regions significant within the 244-node positive-strength regional family, together with additional findings from two other contrast-by-sign families; displayed segments join effect estimates to zero, not confidence intervals. The left cingulate 24b effect also passed the pooled 1,464-test sensitivity correction. Humanized APOE n=261; KO-inclusive n=292; 244 gray-matter nodes. FDR families and model adjustments are as documented in Methods; post-MRI exercise assignment excluded.

### Whole-connectome multivariate distances distinguish humanized APOE groups

The cross-validated connectome-distance test detected genotype-associated multivariate separation among the humanized APOE groups (global permutation p = 0.005; 999 permutations). Displayed cross-validated squared distances per edge (×1,000) were greatest for APOE2 versus KO (0.047), APOE2 versus APOE4 (0.037) and APOE2 versus APOE3 (0.017). APOE3–APOE4 and APOE4–KO estimates were approximately zero or negative and were not represented as meaningful negative metric distances. Despite centroid differences, the PCA showed substantial overlap at the individual-animal level, indicating that genotype-associated group shifts do not imply reliable individual-level discrimination (Figure 2B,C).

### Regional APOE4-associated reductions in positive connectivity center on cingulate area 24b

Of 244 atlas regions examined for each of six genotype-contrast-by-connectivity-sign families, 15 cortical regions showed significantly lower positive nodal strength in APOE4 than APOE2 mice after BH correction within the corresponding 244-region family (all q < 0.05; n = 261; Figure 2H). The largest effect was in left cingulate cortex area 24b (β = −0.653, t = −4.278, two-sided p = 2.69 × 10⁻⁵, within-family q = 0.00656). The significant cortical network encompassed bilateral cingulate areas 24b and 32, right cingulate area 24a′ and left area 29c, bilateral frontal association and primary motor cortex, bilateral forelimb somatosensory cortex, right hindlimb somatosensory cortex, left shoulder somatosensory cortex and right medial orbital cortex. All 15 coefficients were negative, with four additional positive-strength regions showing suggestive within-family q values between 0.05 and 0.10. In the stricter sensitivity analysis applying BH across all 1,464 regional tests, left cingulate area 24b was the only region that remained significant (pooled q = 0.03935). The 15-region cortical pattern is therefore supported within the specified contrast-by-sign family, whereas pooled correction yields a single anatomically localized discovery.

### Apoe knockout comparisons

In the full sample with HN adjustment (31 KO and 261 humanized mice), KO relative to APOE2 was associated with lower positive strength (β = −0.759, 95% CI −1.329 to −0.162; p = 0.0036, q = 0.0324) and clustering (β = −0.628, 95% CI −1.200 to −0.023; p = 0.0084, q = 0.0378). The same comparison showed higher global efficiency (β ≈ +0.59, q = 0.0441) and positive β₁ AUC (β ≈ +0.54, q = 0.0441), while the modularity difference did not reach q < 0.05 (β ≈ −0.44, q = 0.0969). KO–APOE4 comparisons of these six displayed outcomes were not FDR significant. These are absence-of-evidence results rather than formal genotype equivalence tests (Table 1).

### Covariates, risk factors and interaction analyses

The primary genotype associations were adjusted for MRI age, sex, diet, corrected body mass, HN background, protocol and scan year. Separate saved models also evaluate risk factors and genotype-related modifiers, with their own multiplicity families. Because the final numerical outputs of those ancillary models were not included in the archived Figure 2 manuscript document used for this revision, no additional significance claims are made here. Their effect sizes, p values, FDR families and sample sizes should be incorporated only after direct verification against the no-exercise ’risk_factor_tests.csv’ and corresponding interaction-results files; otherwise those analyses remain supporting rather than confirmed main-text findings.

### APOE-dependent cardiac–functional network coupling Global topology: LA × APOE interaction

Among 108 control-diet mice (APOE2 n = 46, APOE3 n = 27, APOE4 n = 35), the association between diastolic left atrial volume and the negative-edge β₀ persistence AUC differed by genotype (joint HC3-robust p = 0.00386; Figure 3A). The interaction survived BH correction across four cardiac exposures (q4 = 0.0154) and across the 12 persistence-topology tests (q12 = 0.0463), but not across all 28 global tests (q28 = 0.108; Table 2).

**Figure 3.**
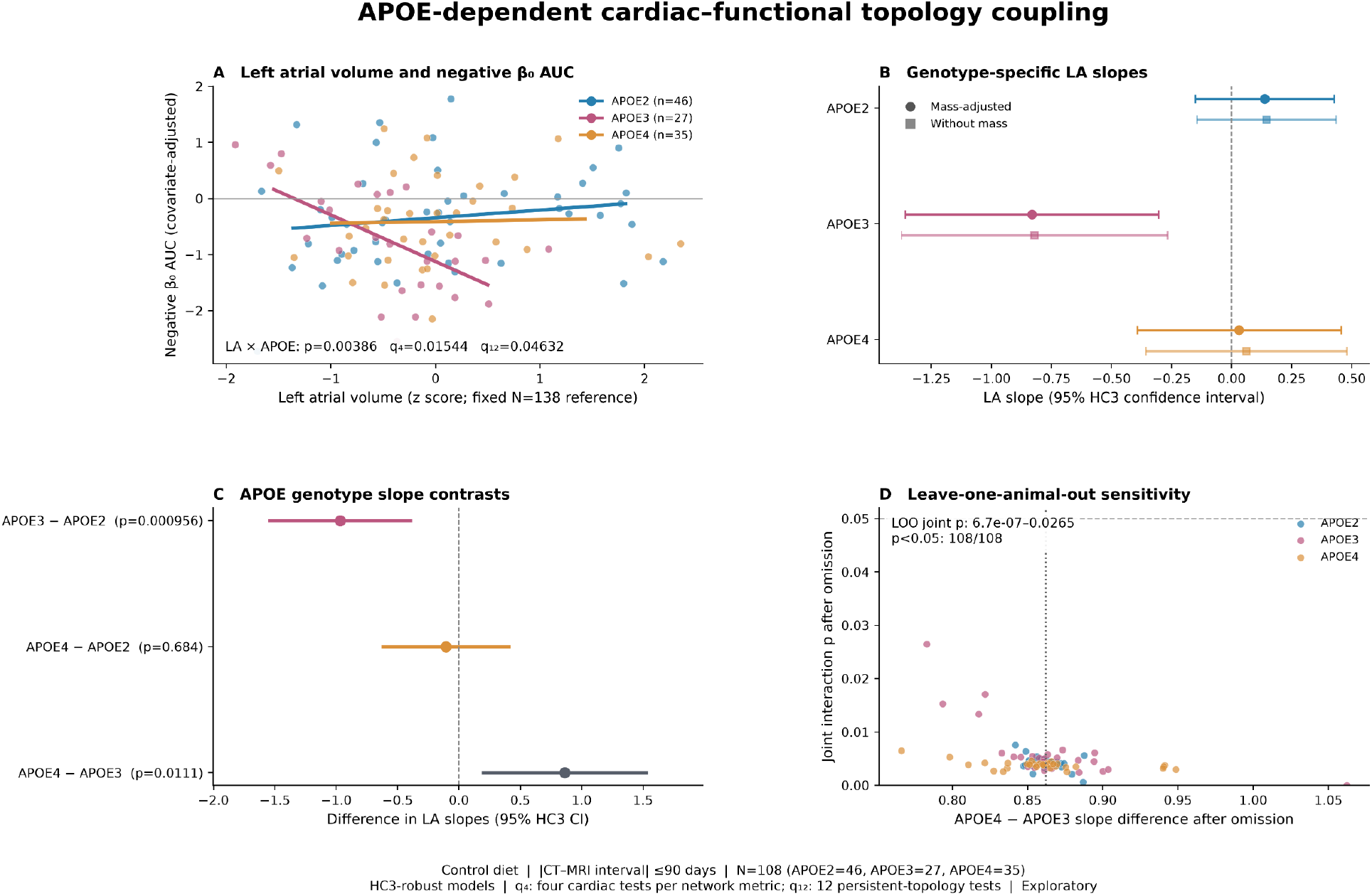
APOE-dependent left atrial volume associations with negative-edge β₀ persistence area under the curve in 108 control-diet humanized APOE mice (APOE2 n = 46; APOE3 n = 27; APOE4 n = 35; absolute CT– MRI interval ≤90 days). (A) Adjusted genotype-specific associations. (B) Genotype-specific slopes and HC3 confidence intervals, with body-mass-adjusted primary and mass-free sensitivity models. (C) Pairwise genotype slope contrasts with HC3 confidence intervals and nominal p values. (D) Leave-one-animal-out sensitivity. The joint two-degree-of-freedom interaction was p = 0.003860318 (q₄ = 0.01544127 within the negative β₀ cardiac family; q₁₂ = 0.04632382 in the persistence domain; q₂₈ = 0.108 across the 28 global tests). All 108 leave-one- out fits retained nominal p < 0.05. The interaction is primarily associated with the negative APOE3 slope and does not establish causality or an APOE4-specific effect.

**Table 2.** Global APOE × left atrial volume interaction on negative β₀ persistence.

| **Analysis** | **N** | **Joint p** | **BH q<sub>4</sub>** | **BH q<sub>12</sub>** | **BH q<sub>28</sub>** |
|---|---|---|---|---|---|
| **Primary, adjusted for mass** | 108 | 0.003860 | 0.015441 | 0.046324 | 0.108 |
| **Without mass** | 108 | 0.005405 | 0.021621 | Not reported for mass-free model | Not calculated |

### Genotype-specific slopes and sensitivity analyses

The genotype-specific LA slope was negative in APOE3 mice (β = −0.831; 95% HC3 CI, −1.358 to −0.304; p = 0.0020) and close to zero in APOE2 (β = 0.138; 95% CI, −0.152 to 0.427) and APOE4 mice (β = 0.031; 95% CI, −0.393 to 0.455; Figure 3B). The APOE3−APOE2 contrast was −0.969 (p = 0.00096), and the APOE4−APOE3 contrast was 0.862 (95% HC3 CI, 0.197 to 1.527; p = 0.0111). The APOE4−APOE2 contrast was −0.106 (p = 0.684; Table 3; Figure 3C). The interaction therefore reflects mainly a negative slope in APOE3 mice rather than a shift specific to APOE4. Removing body mass from the covariates gave a comparable interaction (p = 0.00541; q4 = 0.0216). In leave-one-animal-out analysis, the interaction remained nominally significant in all 108 refits (p range, 6.74 × 10⁻⁷ to 0.0265; Figure 3D). These checks address influence by single animals, not correlated acquisition effects or unmeasured confounding.

**Table 3.** Genotype contrasts for the global LA association.

| **Slope contrast** | **Nominal HC3 p** | **Interpretation** |
|---|---|---|
| **APOE3 − APOE2** | 0.000956 | More negative slope in APOE3 |
| **APOE4 − APOE2** | 0.684 | No reliable difference |
| **APOE4 − APOE3** | 0.0111 | More positive slope in APOE4 than APOE3 |

### Regional APOE × cardiac coupling: primary and expanded exploratory screens

We screened positive and absolute negative nodal connectivity strength in 244 CHASS regions for APOE-by- cardiac interactions in the same 108 control-diet mice. The original four-cardiac-phenotype screen comprised 1,952 tests (488 signed regional outcomes per phenotype). Two interactions passed BH correction within their respective 488-test cardiac families: diastolic/maximal left atrial volume with positive strength in the microcellular tegmental nucleus (p = 6.14 × 10⁻⁵; q₄₈₈ = 0.0299) and diastolic myocardial volume with positive strength in right central gray (p = 4.40 × 10⁻⁵; q₄₈₈ = 0.0215). Neither survived pooled correction across the original 1,952 tests (both q₁₉₅₂ = 0.0599). An expanded exploratory screen of 14 cardiac phenotypes (6,832 tests) additionally identified systolic myocardial volume × right central gray (p = 4.62 × 10⁻⁵; within-family q₄₈₈ = 0.0226), but no regional association survived pooled BH across 6,832 tests (minimum q₆₈₃₂ = 0.1397; Supplementary Figure S3; Supplementary Tables S3A). The original microcellular tegmental and central gray associations retained nominal significance following every single-animal deletion (maximum p = 0.000386 and 0.00244, respectively). In the nested historical subset (n = 80), corresponding nominal joint p values were 0.000828 and 0.00734. These are robustness checks, not independent replication. Analyses restricted post hoc to the largest 20, 40 or 80 CHASS regions yielded no pooled BH discoveries. The regional findings remain hypothesis-generating and are not evidence of mediation of the global negative-edge persistence association (Supplementary Figure S3; Supplementary Table S3 B).

### Spatial navigation and connectivity–behavior associations APOE genotype and water maze acquisition

Across 130 mice, winding decreased over five acquisition days in each APOE group, consistent with improvement in navigation efficiency. Adjusted APOE differences were significant (overall p=2.96 × 10⁻⁵), whereas genotype × day interactions were not supported using either linear-day (p=0.293) or categorical-day (p=0.188) tests (Figure 4A; Table 4). Estimated daily log-winding slopes were −0.201 (APOE2), −0.172 (APOE3), and −0.209 (APOE4). Adjusted winding was higher in APOE4 than APOE2 (β=0.144, BH q=0.00121) and APOE3 (β=0.200, BH q=6.93 × 10⁻⁵), but the slope differences were not statistically significant. These observations support genotype-related overall navigation differences rather than divergent acquisition rates.

**Figure 4.**
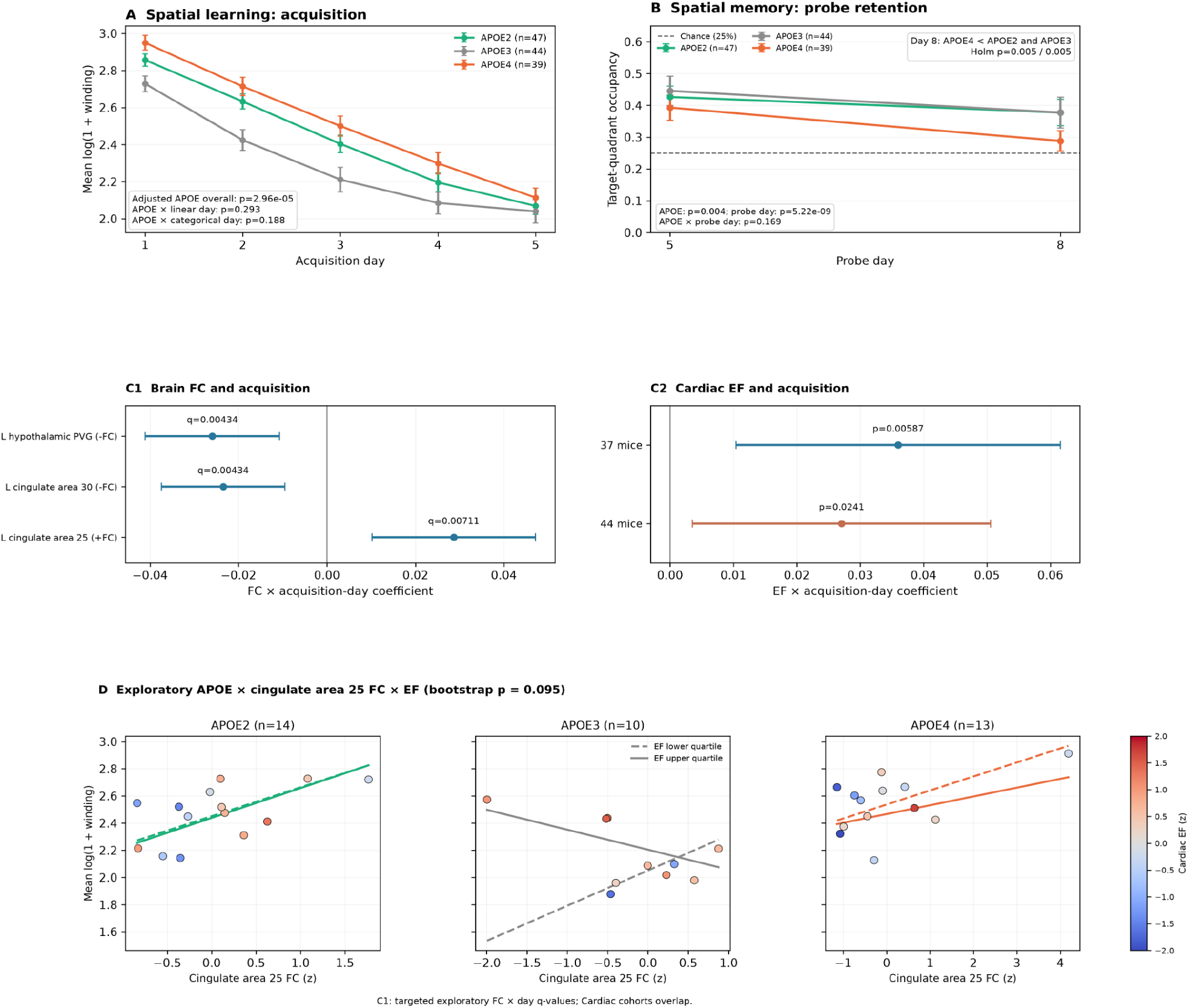
APOE, spatial navigation and brain–heart associations. (A) Five-day acquisition in 130 mice, expressed as log(1+winding), mean ± SEM. (B) Day 5 and Day 8 probe target-quadrant occupancy; dashed line denotes 25% chance, and APOE4 Day 8 occupancy is lower than APOE2 and APOE3 (Holm p=0.00541 each). (C1) Targeted regional FC-by-day coefficients and 95% CIs; only left hypothalamic PVG passes the independent 1,464-test global screen. (C2) EF-by-day associations in primary 37- and overlapping 44-mouse cohorts. (D) Exploratory genotype-stratified cingulate area 25 positive FC and EF analysis; primary wild-cluster bootstrap p=0.095, not significant.

**Table 4.** Principal spatial navigation and neurocardiac behavior results.

| Finding | Mice | Effect / inference | Interpretation |
|---|---|---|---|
| **Acquisition: APOE overall** | 130 | p=2.96×10<sup>−5</sup> | Adjusted genotype effect |
| **Acquisition: APOE × linear day** | 130 | p=0.293 | Interaction not significant |
| **Acquisition: APOE4–APOE2** | 130 | β=0.144; BH q=0.00121 | Adjusted overall winding |
| **Acquisition: APOE4–APOE3** | 130 | β=0.200; BH q=0.0000693 | Adjusted overall winding |
| **Probe: APOE overall** | 130 | p=0.00368 | Paired Day 5/8 analysis |
| **Probe: Day 8–Day 5** | 130 | β=−0.0722; p=5.22×10<sup>−9</sup> | Pooled probe-day effect |
| **Probe: APOE × day** | 130 | p=0.169 | No evidence of different decline |
| **Probe Day 8: APOE4–APOE2** | 130 | β=−0.0875; Holm p=0.00541 | Target-quadrant occupancy |
| **Probe Day 8: APOE4–APOE3** | 130 | β=−0.0886; Holm p=0.00541 | Target-quadrant occupancy |
| **Original unbiased PVG (-FC), Day 5** | 130 | β=−0.1064; global q=0.00501 | Only global 1464-test FDR hit |
| **Targeted FC×day: PVG (-FC)** | 130 | β=−0.0259; targeted q=0.00434 | Different inferential family |
| **Targeted FC×day: cingulate 30 (-FC)** | 130 | β=−0.0235; targeted q=0.00434 | Global screen q=0.0986 |
| **Targeted FC×day: cingulate 25 (+FC)** | 130 | β=0.0287; targeted q=0.00711 | Global screen q=0.0986 |
| **EF × day, primary** | 37 | β=0.0360; p=0.00587 | Within-family BH q=0.0176 |
| **EF × day, sensitivity** | 44 | β=0.0271; p=0.0241 | Within-family BH q=0.0723 |
| **APOE × FC × EF, bootstrap** | 37 | p=0.095 | Exploratory, not significant |

### Paired probe retention distinguishes APOE4 at Day 8

Target-quadrant occupancy decreased between Day 5 and Day 8 across genotypes (adjusted Day 8 minus Day 5 β=−0.0722, p=5.22 × 10⁻⁹). The overall APOE effect was significant (p=0.00368), but the APOE × probe-day interaction was not (p=0.169). On Day 8, APOE4 occupancy was lower than APOE2 (difference −0.0875; Holm p=0.00541) and APOE3 (difference −0.0886; Holm p=0.00541). Consequently, APOE4 exhibited lower delayed probe occupancy, but a statistically greater genotype-specific decline cannot be claimed (Figure 4B; Table 4).

### Regional connectivity relates to acquisition dynamics

In the original unbiased 1,464-test screen, left hypothalamic PVG negative connectivity was associated with Day 5 winding and survived global BH correction (β=−0.1064, q=0.00501). Left cingulate area 30 negative connectivity and left cingulate area 25 positive connectivity had global-screen q=0.0986 and therefore did not reach the 0.05 threshold. Subsequent targeted longitudinal models identified FC × acquisition-day associations for PVG (β=−0.0259; targeted q=0.00434), cingulate area 30 (β=−0.0235; targeted q=0.00434), and cingulate area 25 (β=0.0287; targeted q=0.00711) (Figure 4C1; Table 4). These contrasts represent a different endpoint and a narrower correction family than the unbiased screen.

### Cardiac ejection fraction and acquisition

In mice with adjudicated cardiac–behavior links, standardized EF interacted with acquisition day in the primary 37-mouse cohort (β=0.0360, 95% CI 0.0104 to 0.0616, p=0.00587). The overlapping 44-mouse sensitivity cohort showed a comparable nominal association (β=0.0271, 95% CI 0.00355 to 0.0506, p=0.0241). The latter had within-family BH q=0.0723, whereas the primary EF result had within-family BH q=0.0176 (Figure 4C2). The positive coefficients indicate a less negative change in log-winding across acquisition days at higher EF, conditional on the fitted model; they do not establish a protective or harmful causal effect.

### APOE-dependent brain–heart–behavior patterns remain exploratory

Panel D shows observed mouse-level acquisition-average log-winding versus standardized left cingulate area 25 positive FC, colored by EF, together with model-predicted curves at lower and upper EF quartiles in APOE2 (n=14), APOE3 (n=10), and APOE4 (n=13). Although the trial-cluster asymptotic joint test suggested a strong interaction (p≈2.27 × 10⁻⁶), conservative analyses did not validate it: the primary trial-level wild-cluster bootstrap gave p=0.095 (mouse-day bootstrap p=0.090; mouse-level HC3 p=0.762). The 44-mouse overlapping sensitivity analysis similarly gave bootstrap p=0.080–0.0875. Within-genotype contrasts showed the most pronounced nominal FC × EF pattern in APOE3 (bootstrap p=0.0825), but none of six contrasts survived Holm adjustment (minimum adjusted p=0.495). The joint model including EF and cingulate FC also yielded no EF/FC effects surviving its two-test Holm adjustment (Supplementary Table S4).

### Molecular context of APOE-associated network effects

#### A nominal excitatory-neuron spatial correspondence with the APOE4 network map

Regions with stronger excitatory-neuron molecular signatures tended to show more negative APOE4−APOE2 effects on positive nodal strength. Under hand mapping, mean gene-wise Spearman r was −0.4051 (spatial-null p = 0.0292; random-gene-set p = 0.0158; combined p = 0.0292; FDR q = 0.2044). Registration-based mapping gave r = −0.3911 (spatial-null p = 0.0442; random-set p = 0.0208; combined p = 0.0442; FDR q = 0.3093). The directional pattern persisted across atlas mappings but neither test passed the specified gene-set FDR correction. No cell-type or pathway test in the 32-row original mapping-specific family met q < 0.05 (Supplementary Table S5). These genotype-associated molecular signatures and their adjusted contrasts are illustrated in Figure 5.

**Figure 5.**
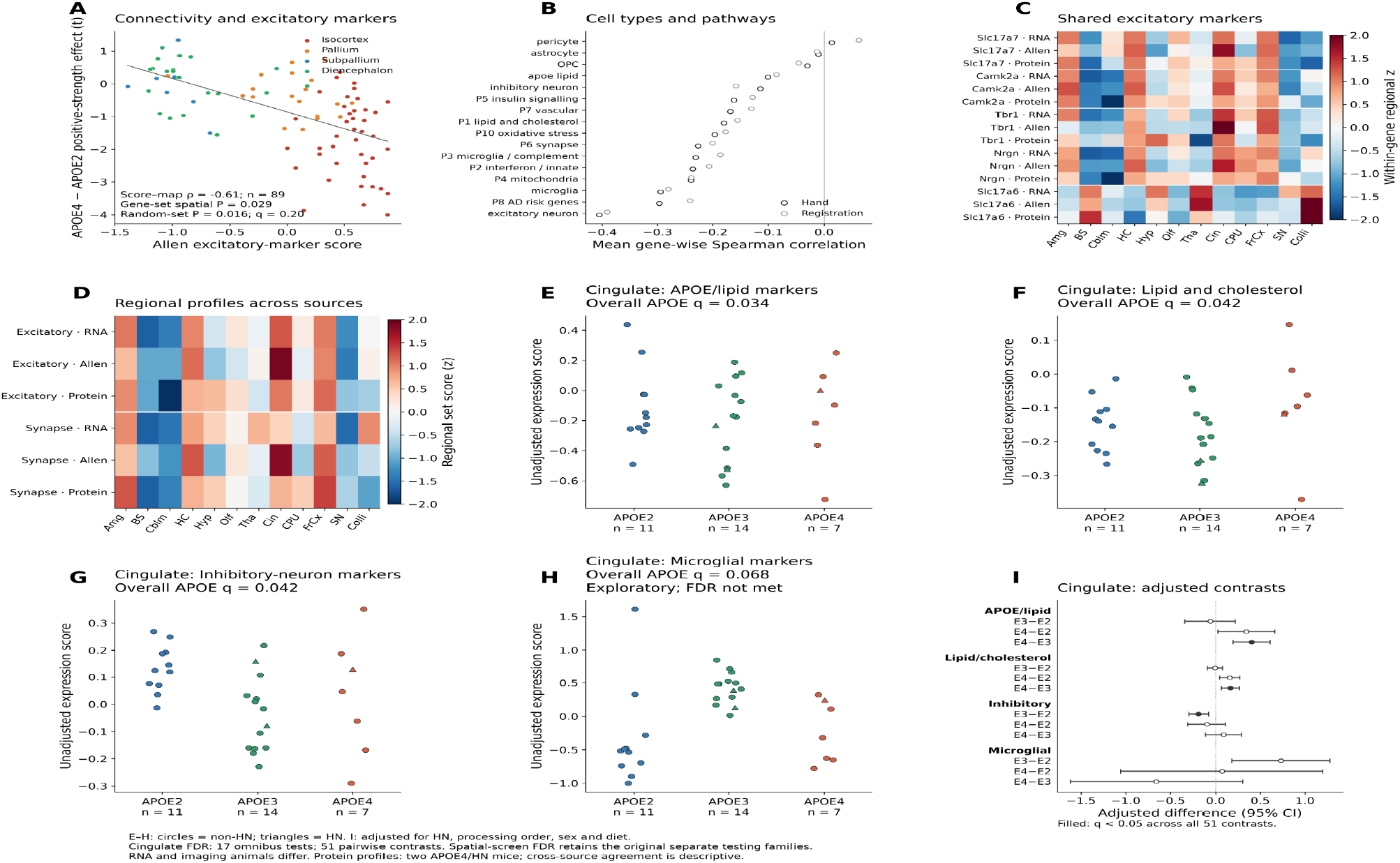
Molecular context of APOE-associated functional-network differences and cingulate gene-expression signatures. (A) Allen excitatory-neuron marker score against the APOE4–APOE2 positive nodal-strength effect across 89 mapped structures, colored by anatomical division. The displayed score-map correlation is distinct from the gene-wise spatial-null inference (spatial p = 0.029, random- gene-set p = 0.016; gene-set FDR q = 0.204). (B) Mean gene-wise Spearman associations between cell-type/pathway scores and the imaging effect map using hand and registration-based atlas correspondences; no gene set passed FDR correction. (C) Regional heat maps of five shared excitatory-neuron markers across laboratory RNA-seq, Allen reference expression and exploratory proteomics, expressed as within-gene regional z scores. (D) Regional excitatory-neuron and synapse gene-set profiles across these three sources. Cross-platform concordance describes anatomical expression patterns, not replicated genotype effects. (E–H) Individual-animal cingulate APOE/lipid-marker, lipid/cholesterol, inhibitory-neuron and microglial-marker scores, respectively, stratified by APOE genotype. Circles denote HN-negative and triangles HN-positive mice. Overall APOE BH q values across 17 prespecified signatures are 0.0336, 0.0419, 0.0419 and 0.0677, respectively; the microglial result did not pass FDR. (I) Adjusted pairwise genotype contrasts and nominal 95% HC3 confidence intervals; filled points indicate BH q < 0.05 over all 51 planned contrasts. The cingulate sample includes 32 mice (APOE2 n = 11, APOE3 n = 14, APOE4 n = 7). Models adjusted for HN status, processing order, sex and diet. Imaging and RNA-seq animals were distinct; proteomics from two APOE4/HN mice was descriptive.

#### Concordant regional neuronal expression across platforms

Across 12 dissected regions, excitatory-marker scores correlated between laboratory RNA-seq and Allen expression (r = 0.881), RNA-seq and proteomics (r = 0.895), and Allen and proteomics (r = 0.804) (Table 5). Corresponding correlations for the synapse GO set were 0.559, 0.573 and 0.720. These findings support the anatomical interpretability of excitatory-neuron enrichment across resources, not molecular replication of APOE genotype effects. Protein measures were descriptive averages from only two APOE4/HN mice (Supplementary

**Table 5.** Key molecular findings.

| **Analysis** | **Effect / sample** | **Multiplicity / interpretation** |
|---|---|---|
| **Allen excitatory mapping, hand** | Mean gene-wise r=−0.4051; spatial p=0.0292 | BH q=0.2044; not FDR-significant |
| **Allen excitatory mapping, registration** | Mean gene-wise r=−0.3911; spatial p=0.0442 | BH q=0.3093; not FDR-significant |
| **Cingulate APOE/lipid omnibus** | F(2,25)=8.069; n=32 | BH q=0.0336 (17 tests) |
| **Cingulate inhibitory markers omnibus** | F(2,25)=6.355; n=32 | BH q=0.0419 (17 tests) |
| **Cingulate lipid/cholesterol omnibus** | F(2,25)=6.010; n=32 | BH q=0.0419 (17 tests) |
| **APOE4 vs APOE3, APOE/lipid** | β=+0.4023 | BH q=0.0245 (51 contrasts) |
| **APOE4 vs APOE3, lipid/cholesterol** | β=+0.1649 | BH q=0.0395 (51 contrasts) |
| **APOE3 vs APOE2, inhibitory markers** | β=−0.1844 | BH q=0.0390 (51 contrasts) |

Table S5). The concordance of regional excitatory-neuron and synaptic signatures across laboratory RNA-seq, Allen reference expression, and exploratory proteomics supports the anatomical consistency of these molecular profiles, although the two-animal proteomics dataset does not permit inference about APOE genotype effects.

#### Three corrected molecular signatures in cingulate cortex

Three of 17 predefined cingulate scores showed adjusted omnibus APOE differences after FDR correction: APOE/lipid markers (F(2,25) = 8.069, p = 0.00198, q = 0.0336); inhibitory-neuron markers (F(2,25) = 6.355, p = 0.00587, q = 0.0419); and lipid/cholesterol processes (F(2,25) = 6.010, p = 0.00739, q = 0.0419). Across the 51 planned contrasts, APOE4 mice had higher APOE/lipid-marker scores than APOE3 (+0.4023; 95% CI [0.1958, 0.6089]; q = 0.0245) and higher lipid/cholesterol scores (+0.1649; [0.0647, 0.2651]; q = 0.0395), while APOE3 had lower inhibitory-marker scores than APOE2 (−0.1844; [−0.2911, −0.0776]; q = 0.0390). Microglial markers were nominally different by genotype (p = 0.0159) but did not pass FDR (q = 0.0677).

In 29 non-HN mice, the three corrected contrasts retained the same sign (APOE/lipid +0.4119, p = 0.00486; lipid/cholesterol +0.1628, p = 0.0470; inhibitory −0.1842, p = 0.00195). No sign changes occurred across any of the 32 leave-one-animal-out fits per contrast (Supplementary Tables F5-S5 and F5-S6). The small APOE4 subgroup and sequencing-order imbalance nevertheless constrain causal or mechanistic interpretation.

### APOE genotype modifies regional brain MRI–cardiac associations

We examined APOE-dependent associations between cardiac CT phenotypes and regional brain volume, intrinsic T1, and cerebral blood flow (CBF) in modality-specific cohorts (volume, n = 92; T1 and CBF, n = 64 each). The HN-harmonized APOE-by-cardiac interaction models included age, sex, exercise assignment, and humanized NOS2 status, with diet included when it varied within the complete-case cohort. Benjamini–Hochberg false-discovery-rate (BH-FDR) correction was applied separately to the planned region-by-cardiac families (4,326 volume tests and 4,312 tests each for T1 and CBF). Significant interactions comprised 12 volume associations across three CHASS regions, 93 T1 associations across 61 regions, and 36 CBF associations across 29 regions (Figure 6A–I; Table 6; Supplementary Figure S6).

**Figure 6.**
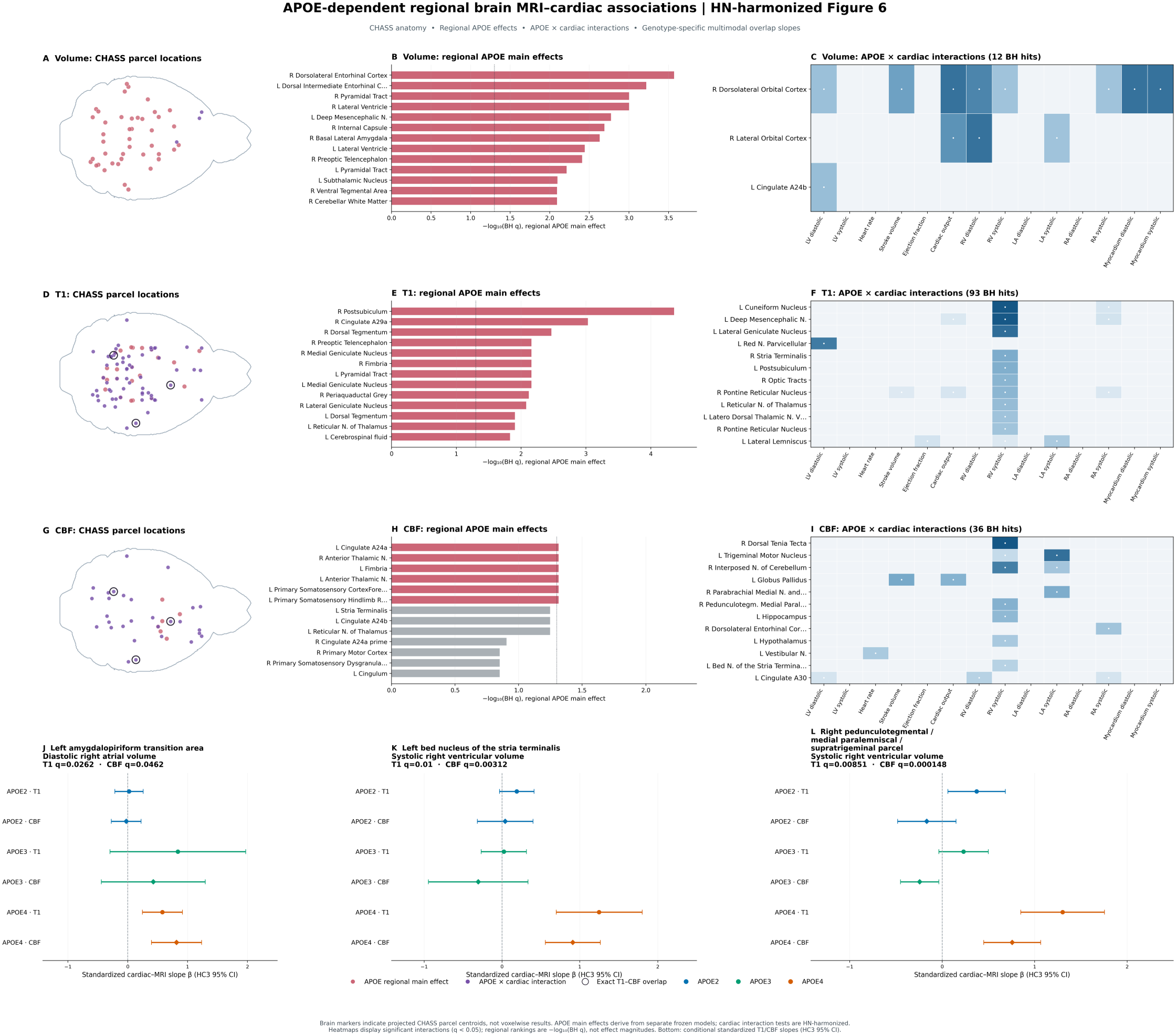
APOE genotype modifies associations between regional brain MRI phenotypes and cardiac CT measurements. Rows 1–3 present volume, intrinsic T1, and CBF, respectively. Panels A, D, and G display CHASS brain silhouettes with region-centroid markers for significant APOE main effects (rose) and APOE-by- cardiac interactions (purple); circled markers denote regions contributing to an exact T1–CBF overlap. Panels B, E, and H rank FDR-significant regional APOE main effects. Panels C, F, and I summarize significant region– cardiac interaction tests as heatmaps. Panels J–L present standardized genotype-specific slopes (APOE2, blue; APOE3, green; APOE4, vermilion) and HC3 95% confidence intervals for the three exact T1–CBF region–cardiac associations: left amygdalopiriform transition area with diastolic right atrial volume, left BNST with systolic right ventricular volume, and right pedunculotegmental/medial paralemniscal/supratrigeminal composite region with systolic right ventricular volume. Reported q values are modality-family BH-FDR-adjusted omnibus APOE-by- cardiac interaction values. Centroid markers localize CHASS parcels and do not represent voxelwise statistical effects. The regional main-effect models were fitted separately from the APOE-by-cardiac interactions.

**Table 6.**
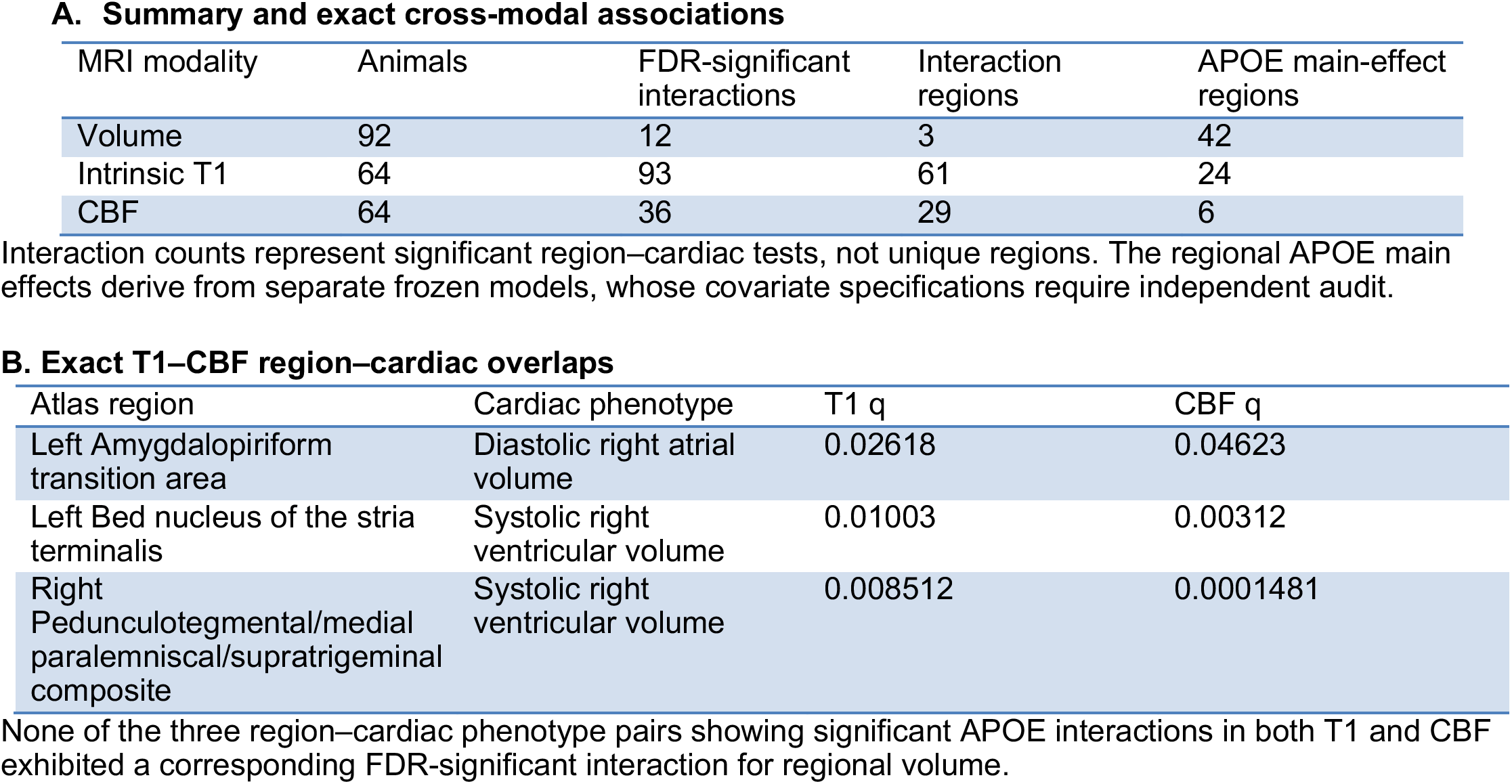
Cross Modal Associations.

| A. Summary and exact cross-modal associations |  |  |  |  |
|---|---|---|---|---|
| MRI modality | Animals | FDR-significant interactions | Interaction regions | APOE main-effect regions |
| Volume | 92 | 12 | 3 | 42 |
| Intrinsic T1 | 64 | 93 | 61 | 24 |
| CBF | 64 | 36 | 29 | 6 |
| **B. Exact T1–CBF region–cardiac overlaps** |  |  |  |  |
| Atlas region | Cardiac phenotype | T1 q | CBF q |  |
| Left Amygdalopiriform transition area | Diastolic right atrial volume | 0.02618 | 0.04623 |  |
| Left Bed nucleus of the stria terminalis | Systolic right ventricular volume | 0.01003 | 0.00312 |  |
| Right Pedunculotegmental/medial paralemniscal/supratrigeminal composite | Systolic right ventricular volume | 0.008512 | 0.0001481 |  |

Interaction counts represent significant region–cardiac tests, not unique regions. The regional APOE main effects derive from separate frozen models, whose covariate specifications require independent audit.
None of the three region–cardiac phenotype pairs showing significant APOE interactions in both T1 and CBF exhibited a corresponding FDR-significant interaction for regional volume.

Volume–cardiac interactions were concentrated in the right dorsolateral orbital cortex (eight associations) and right lateral orbital cortex (three associations), with one additional association between left cingulate cortex area 24b and diastolic left ventricular volume (q = 0.0275). Right dorsolateral orbital cortex interactions included cardiac output (q = 0.00223), stroke volume (q = 0.00828), and additional myocardial, atrial, and ventricular phenotypes. Right lateral orbital cortex interactions involved cardiac output (q = 0.00558), diastolic right ventricular volume (q = 0.00223), and systolic left atrial volume (q = 0.0336; Supplementary Table S6A).

Three exact CHASS region–cardiac phenotype pairs exhibited FDR-significant APOE interactions in both T1 and CBF (Figure 6J–L; Table 6). Diastolic right atrial volume was associated with left amygdalopiriform transition area T1 (q = 0.0262) and CBF (q = 0.0462). Systolic right ventricular volume was associated with T1 and CBF in the left bed nucleus of the stria terminalis (BNST; q = 0.0100 and 0.00312, respectively) and the right pedunculotegmental/medial paralemniscal/supratrigeminal composite parcel (q = 0.00851 and 0.000148, respectively). No exact region–cardiac pair met FDR significance for regional volume in addition to T1 or CBF.

For all three multimodal pairs, APOE4-specific standardized slopes were positive for both T1 and CBF, with heteroskedasticity-consistent (HC3) 95% confidence intervals excluding zero. In the left amygdalopiriform transition area, APOE4 slopes were β = 0.580 (95% CI 0.246–0.913) for T1 and β = 0.815 (0.396–1.234) for CBF. Corresponding T1 and CBF slopes were 1.249 (0.698–1.799) and 0.911 (0.557–1.266) for BNST, and 1.302 (0.849–1.755) and 0.757 (0.449–1.065) for the pontine composite parcel. These slopes describe conditional associations; the interaction q values quantify the omnibus two-degree-of-freedom genotype-by- cardiac test, not the significance of individual slopes. None of the three exact multimodal associations coincided with a significant regional APOE main effect for its T1 or CBF measurement (Figure 6J–L; Table 6; Supplementary Figure S7; Supplementary Table S6 B-C).

Separate regional APOE main-effect analyses yielded 42 volume, 24 T1, and six CBF significant regions (Figure 6B, E, H; Supplementary Table S6D). Eight T1 regions (nine region–cardiac pairs) and one CBF region (one pair) showed both a significant regional main effect and at least one genotype-dependent cardiac association; no volume region showed both. Right cingulate cortex area 29a had a T1 APOE main effect (q = 0.000931) and an APOE-by-cardiac output interaction (q = 0.0262); left fimbria showed a CBF main effect (q = 0.0484) and an interaction involving diastolic right ventricular volume (q = 0.0205). These patterns distinguish differences in mean MRI measures from genotype-dependent brain–cardiac covariation (Supplementary Table S6D).

The cross-modal observations represent convergence of statistical associations within common anatomical parcels, not evidence of a causal neurocardiac pathway. Interpretation is limited by the interval between MRI and CT, small APOE4 representation in the 64-animal cohorts, and sensitivity to covariate specification. CT- associated mass was not included in the selected primary analysis; the separate M4 + HN sensitivity analysis gave 8, 86, and 33 FDR-significant interactions for volume, T1, and CBF, respectively, and retained one exact T1–CBF overlap.

## Discussion

### APOE isoforms shape functional brain networks and their cardiovascular associations

We investigated whether APOE isoforms influence functional brain network organization and modify associations between cerebral and cardiovascular phenotypes. The principal finding was a distinct APOE4-associated functional network architecture, characterized by reduced positive connectivity strength, clustering, and modularity, together with greater global efficiency and altered persistent topology. These differences were accompanied by regional reductions in positive connectivity involving predominantly cortical regions, particularly the cingulate cortex. In mice with paired cardiac CT and resting-state fMRI, the association between left atrial volume and negative-network β₀ persistence differed by APOE genotype, primarily reflecting a negative association in APOE3 mice. Complementary analyses identified genotype-dependent associations between regional cardiac phenotypes and brain volume, intrinsic T1, or cerebral blood flow (CBF), including anatomically matched T1–CBF findings in limbic-related and pontine regions. Behavioral differences and independent cingulate transcriptomic signatures provided additional biological context. Together, these findings suggest that APOE-dependent variation extends beyond individual brain or cardiac phenotypes to their observed relationships, although the direction and mechanisms of these associations remain unresolved.

### APOE-dependent reorganization of functional brain networks

The reduced positive network strength, clustering, and modularity in APOE4 relative to APOE2 mice indicate differences in large-scale functional network organization. These findings are consistent with previous reports of altered structural and functional connectivity in humanized APOE mouse models (Badea et al., 2019; Zerbi et al., 2014; Winter et al., 2024). The concurrent increase in global efficiency and changes in persistent topology suggest a different balance between local network segregation and graph-wide integration. However, greater graph efficiency should not be interpreted as improved neural information processing, particularly when accompanied by weaker positive connectivity and reduced clustering. Similarly, persistence measures characterize topological features across network thresholds rather than identifiable physiological feedback circuits.

The regional findings provide anatomical specificity. Fifteen cortical regions exhibited reduced positive nodal strength in APOE4 relative to APOE2 mice within the designated 244-region correction family, with left cingulate area 24b remaining significant after correction across all 1,464 regional tests. The affected regions included additional cingulate, frontal association, motor, and somatosensory territories. This distribution suggests that APOE-associated network differences extend across distributed cortical systems rather than being confined to regions traditionally implicated in memory. Because resting-state functional connectivity reflects coordinated hemodynamic fluctuations, these findings may arise from neuronal network differences, altered neurovascular regulation, or contributions from both processes.

The *Apoe*-knockout animals provided an additional perspective on isoform-associated network variation. Relative to APOE2, knockout mice exhibited lower positive connectivity strength and clustering, together with greater global efficiency and positive β₁ persistence, partly resembling the APOE4 phenotype. Previous resting- state fMRI studies have identified connectivity differences in APOE4 and *Apoe*-null mice (Zerbi et al. 2014). Nevertheless, the lack of significant APOE4-versus-knockout differences does not establish functional equivalence or indicate that APOE4 acts through a simple loss-of-function mechanism. The restricted acquisition protocol and HN background of the knockout cohort further limit mechanistic comparisons.

### APOE modifies associations between cardiac phenotypes and functional network topology

A central contribution of this study is the identification of APOE-dependent associations between cardiac structure and functional network topology. In the 108-mouse paired cohort, left atrial volume showed a genotype interaction with negative-network β₀ persistence (joint p = 0.00386), surviving FDR correction within the four- cardiac-exposure family (q = 0.0154) and the 12-test persistence family (q = 0.0463), but not across all 28 global tests (q = 0.108). This distinction is important given the exploratory selection of cardiac predictors. The interaction was stable to individual-animal removal and body-mass adjustment, but these sensitivity analyses do not address all sources of residual confounding. Notably, the association was driven predominantly by a negative left atrial volume–persistence slope in APOE3 mice, rather than an accentuated effect in APOE4. This finding argues against assuming a monotonic relationship between Alzheimer’s disease risk and every cardiac–neural association. APOE isoforms may influence the structure of cross-organ relationships in qualitatively different ways, rather than simply increasing or decreasing the magnitude of a common effect.

The physiological interpretation requires caution. Left atrial volume reflects cardiac structure and loading conditions, whereas negative-network β₀ persistence describes the evolution of connected components across a functional-connectivity filtration (Stolz, Harrington, and Porter 2017) (Songdechakraiwut and Chung 2020). Neither measurement directly quantifies cardiac autonomic drive, neural firing, or synaptic inhibition. The association may reflect neural influences on cardiovascular regulation, afferent cardiac signaling affecting brain activity, or shared APOE-dependent metabolic, vascular, inflammatory, and developmental processes. Moreover, the noncontemporaneous MRI and CT acquisitions preclude inference about temporal direction.

Regional connectivity–cardiac analyses identified candidate associations involving the microcellular tegmental nucleus and right central gray. Although selected interactions were significant within individual cardiac-specific testing families and showed sensitivity stability, none survived pooled FDR correction across the expanded 6,832-test regional screen. These findings therefore provide anatomical hypotheses rather than confirmed regional mediators of the global network association.

### Relation to human brain–heart imaging and cardiovascular genetics

The present findings complement human studies demonstrating relationships between cardiac structure or performance and cerebral morphology, microstructure, and functional connectivity. In a large multiorgan imaging analysis, Zhao et al. (Zhao et al. 2023) identified shared phenotypic and genetic associations between cardiac and brain MRI traits, suggesting that common genetic factors contribute to variation across these organs. Additional imaging-genetic studies have extended this framework to broader multiorgan relationships (Shu et al. 2025; Zalesky et al. 2026). More specifically, Moore et al. (Moore et al. 2026) reported that lower cardiac output was associated with greater longitudinal brain atrophy among APOE-ε4 carriers, providing human evidence that APOE may modify associations between cardiovascular performance and cerebral structure. Although that study examined structural brain outcomes and longitudinal cardiac relationships rather than resting-state functional topology, it supports the translational relevance of evaluating cardiac–brain associations within genetically defined subgroups.

Our experimental model offers a complementary perspective by comparing three human APOE isoforms and examining functional networks together with quantitative cardiac phenotypes. Previous photon-counting CT studies in humanized APOE mice have demonstrated genotype-, sex-, diet-, and immune-background- dependent differences in cardiac anatomy and responses to exercise(Allphin et al. 2023; Allphin et al. 2025) (Nadkarni et al. 2025). The present study extends this work by investigating whether cardiac variation is associated with distinct neural phenotypes across APOE genotypes. These findings identify measurable outcomes for future experimental tests of brain–heart interactions.

### Regional MRI–cardiac relationships reveal multimodal anatomical convergence

The regional MRI analyses provide complementary evidence that APOE modifies associations between cerebral characteristics and cardiac phenotypes. Using the HN-harmonized models, significant APOE-by-cardiac interactions were detected for normalized regional brain volume, intrinsic T1, and CBF. These interactions were not synonymous with mean differences in regional MRI measurements. Only eight T1 parcels and one CBF parcel showed both a regional APOE main effect and a genotype-dependent cardiac association, with no corresponding overlap among volume parcels. Thus, APOE-dependent covariation between brain and cardiac traits may be detectable even when genotype-related mean differences in the regional MRI measure are not.

Three anatomically matched region–cardiac associations were significant in both intrinsic T1 and CBF: the left amygdalopiriform transition area with diastolic right atrial volume, the left bed nucleus of the stria terminalis (BNST) with systolic right ventricular volume, and a right pedunculotegmental/medial paralemniscal/supratrigeminal composite parcel with systolic right ventricular volume. These associations involved positive APOE4-specific slopes in both modalities, although the statistical evidence for genotype dependence derives from omnibus interaction tests rather than the individual slopes. The matched findings did not extend to regional volume and were not accompanied by significant regional APOE main effects in the corresponding T1 or CBF measurements. The BNST is particularly relevant because it integrates limbic, autonomic, and neuroendocrine responses through connections with hypothalamic and brainstem regulatory systems. Experimental studies have implicated BNST neurotransmission in the modulation of blood pressure, heart rate, baroreflex responses, and stress-related cardiovascular regulation (Crestani et al. 2013). The amygdalopiriform transition area is likewise associated with limbic circuitry, although its contribution to cardiovascular control is less established. The right pontine composite parcel includes heterogeneous tegmental and orofacial sensorimotor structures and cannot be interpreted as a single cardiovascular-regulatory nucleus. Together, these anatomical relationships provide biological plausibility but not evidence that the measured MRI associations arise from direct autonomic pathways. Intrinsic T1 reflects multiple tissue properties, while CBF reflects vascular supply and regulation; both may vary with systemic physiology independently of neural activity. Furthermore, T1 and CBF were measured in overlapping animals, making their agreement multimodal anatomical convergence rather than independent replication. The regional associations also differed from the left atrial–network persistence result. Whereas the global functional topology analysis emphasized left atrial volume, the regional MRI analyses involved right atrial and ventricular volumes, cardiac output, and other cardiac phenotypes. Correlation among cardiac measurements and differences in analytical cohorts preclude identification of a single dominant cardiac determinant.

### Cingulate cortex as a convergent anatomical system

The cingulate cortex emerged as a recurring anatomical feature across functional connectivity, regional MRI, and independent molecular analyses. APOE4-associated reductions in positive nodal strength were particularly pronounced in left cingulate area 24b, while the regional MRI findings implicated additional cingulate subdivisions, including right area 29a in T1-related analyses. Independent cingulate RNA sequencing identified APOE-associated differences in lipid/cholesterol-related and inhibitory-neuron marker signatures. These observations suggest convergence at the broader cingulate-system level, although they do not represent identical anatomical parcels or measurements obtained in the same animals. The physiological relevance of the cingulate cortex is supported by its involvement in autonomic and cardiovascular regulation. Human neuroimaging and lesion evidence has linked anterior cingulate activity to autonomic modulation of cardiac function, including sympathetic influences on heart rate (Critchley et al. 2003). Cortical autonomic networks interact with insular, hypothalamic, and brainstem systems that integrate interoceptive information and regulate cardiovascular responses (Cechetto 2014) (Valenza, Matić, and Catrambone 2025). The cingulate network phenotype may therefore be relevant to broader systems coordinating behavioral, affective, and cardiovascular regulation. Nevertheless, the cingulate findings must be interpreted anatomically and statistically. The functional- connectivity effect was localized most strongly to area 24b, whereas regional T1 and CBF findings involved other subdivisions, and the RNA-seq samples represented a larger dissected cingulate territory. Consequently, these results support convergence within a cortical system rather than demonstrating a specific cingulate-to-heart circuit. The independent molecular cohort also cannot establish that lipid or inhibitory-signaling changes mediate the observed functional-network or cardiac associations.

### Neurovascular and tissue-sensitive mechanisms

APOE-dependent vascular biology offers an alternative and potentially complementary explanation for the functional network findings. APOE4 and *Apoe* deficiency have been linked to cerebrovascular injury involving pericyte-associated inflammatory signaling and blood–brain barrier disruption (Bell et al. 2012). Human studies have likewise identified APOE4-associated blood–brain barrier dysfunction relevant to cognitive decline (Montagne et al. 2020). These observations suggest that APOE genotype may alter cerebral vascular regulation independently of—or in interaction with—neuronal network organization. This distinction is particularly important for resting-state BOLD functional connectivity. Systemic hemodynamic variations, regional perfusion delays, and local cerebrovascular properties can influence correlations among BOLD signals. (Schneider et al. 2023) demonstrated that systemic perfusion-related factors may affect resting-state functional connectivity, emphasizing the need to distinguish hemodynamic from neural contributions. Thus, some of the APOE- associated network differences observed here may reflect differences in the vascular transfer of neural activity into BOLD signals rather than altered neuronal communication alone.

Regional CBF measurements provide complementary physiological information but do not directly resolve this ambiguity. Our pCASL approach has previously been applied to multimodal MRI characterization in CVN-AD mice (Badea et al. 2025). Intrinsic T1 provides an additional quantitative, tissue-sensitive measure, but its interpretation is nonspecific because relaxation depends on tissue water, macromolecular composition, and other microstructural properties (Saltarelli et al. 2025). Because functional connectivity, perfusion, intrinsic T1, and cardiac physiology were not jointly evaluated as mediators within a single fully paired longitudinal model, the present findings cannot distinguish neuronal network changes from cerebrovascular influences or shared systemic processes. This limitation is central to the biological interpretation of the multimodal results.

### Behavioral and molecular relevance of APOE-dependent networks

Behavioral analyses demonstrated APOE-associated differences in spatial navigation and delayed memory performance. APOE4 mice exhibited higher absolute winding number during Morris water maze acquisition and lower target-quadrant occupancy in the delayed probe test than APOE2 or APOE3 mice. However, genotype- by-acquisition-day interactions were not significant, indicating differences in overall navigation performance rather than different learning rates. Absolute winding number provides complementary information about spatial search trajectories beyond conventional measures such as escape latency or swimming distance (Badea et al. 2022). Regional functional connectivity was associated with selected navigation measures, including an association between left hypothalamic PVG negative connectivity and Day 5 absolute winding number that survived correction across the unbiased regional screen. Targeted cingulate connectivity analyses provided additional associations with acquisition trajectories, but these did not represent independent discoveries in the full regional testing family. Cardiac ejection fraction was also associated with acquisition trajectories. Higher ejection fraction was associated with a less negative change in log-transformed absolute winding number across acquisition days. This finding indicates a relationship between cardiac performance and navigation dynamics but does not establish a beneficial or adverse causal effect. Moreover, the exploratory APOE-by-cingulate- connectivity-by-ejection-fraction interaction was not supported by conservative bootstrap inference, limiting evidence for a unified genotype-dependent brain–heart–behavior pathway.

Independent transcriptomic analyses provided complementary molecular context. In cingulate cortex, APOE genotype was associated with lipid/cholesterol-related and inhibitory-neuron marker signatures after correction for multiple testing. These findings are compatible with established roles of APOE in lipid transport, glial biology, and neuroimmune regulation (Belloy, Napolioni, and Greicius 2019) (Yamazaki et al. 2019) (Blumenfeld et al. 2024). However, the transcriptomic data were obtained in different animals from the imaging cohort, and sequencing-order imbalance limits interpretation of some genotype contrasts. Spatial correlations between regional functional connectivity effects and excitatory-neuron expression profiles did not survive gene-set FDR correction. Accordingly, the molecular findings should be considered biological context rather than evidence of a demonstrated molecular mechanism linking APOE genotype, functional networks, and cardiac physiology.

### Strengths and limitations

The principal strengths of this study include the genetically defined humanized APOE models, the large functional connectome cohort, complementary graph-theoretical and persistent-homology analyses, and the inclusion of paired cardiac CT–functional MRI measurements. The regional MRI analyses extend the characterization of brain–cardiac relationships beyond resting-state connectivity, while behavioral and molecular data provide complementary functional and biological perspectives. Explicit multiple-testing correction, cohort- specific eligibility criteria, and sensitivity analyses strengthen interpretation of the principal statistical findings.

Several limitations warrant emphasis. First, most measurements were observational and acquired at different time points, preventing inference about the direction of brain–heart influences. Second, although genotype and major biological and acquisition covariates were included in the regression models, residual confounding by anesthesia, scanner protocol, acquisition year, systemic physiology, and experimental exposures remains possible. Third, the regional T1 and CBF cohorts contained only eight APOE4 mice, making genotype-specific slope estimates particularly vulnerable to influential observations. Selected regional findings were also sensitive to HN adjustment, underscoring the importance of model harmonization.

Fourth, the significance of the left atrial–negative-network persistence interaction depended on the multiple- testing family: it survived correction within the persistence domain but not across all global cardiac–network tests. Likewise, regional functional connectivity–cardiac associations did not survive pooled correction across the expanded screening family. Fifth, multimodal convergence frequently involved different anatomical subdivisions, and partially overlapping cohorts do not constitute independent replication. Finally, independent transcriptomic measurements and exploratory behavioral associations cannot establish molecular mediation or a causal neural pathway linking cortical network organization with cardiac function.

### Translational implications and future directions

The present findings support a model in which APOE isoforms influence brain network organization and modify observed associations between neural and cardiovascular phenotypes. The combination of functional connectivity, persistent topology, quantitative regional MRI, and cardiac CT provides candidate imaging outcomes for future studies of APOE-sensitive brain–heart interactions. Their translational relevance is strengthened by emerging human evidence that APOE genotype modifies associations between cardiovascular function and cerebral structure (Moore et al. 2026), although these candidate measures have not been validated for individual risk prediction.

Future studies should prioritize simultaneous or longitudinal assessment of cerebral network dynamics, CBF, cardiac physiology, and autonomic function in the same APOE-defined animals. Measurements of heart-rate variability, arterial pressure, and baroreflex responses would provide more direct autonomic phenotypes than cardiac chamber volumes alone. Targeted perturbation of candidate cingulate–hypothalamic–brainstem pathways, coupled to cardiovascular readouts, could distinguish efferent neural regulation from cardiovascular feedback or shared systemic mechanisms. Interventions influencing APOE-dependent metabolic and vascular pathways may further clarify whether network and cardiac phenotypes respond through common or separable mechanisms.

## Conclusions

Human APOE isoforms are associated with distinct functional brain network architectures and genotype- dependent relationships between cerebral and cardiac phenotypes. APOE4 mice exhibited reduced positive network strength and segregation, with prominent cortical connectivity differences involving the cingulate system. The principal paired brain–heart association linked left atrial volume to negative-network persistence in an APOE-dependent manner, driven predominantly by APOE3. Complementary regional MRI findings identified genotype-dependent T1–CBF–cardiac associations in limbic-related and pontine regions, while behavioral and independent cingulate molecular analyses provided additional biological context. These findings establish a multimodal framework for investigating APOE-sensitive brain–heart relationships but do not determine their causal direction or underlying mechanisms.

## Acknowledgments and funding

This work was supported by the Duke University School of Medicine and the National Institutes of Health under awards RF1 AG057895, R01 AG066184, R56 AG066184, R01 AG070149, R01 CA302738, and R01 HL179758.

## Notes

### Competing Interest Statement

The authors have declared no competing interest.

## References

Allphin, A. J., A. Mahzarnia, D. P. Clark, Y. Qi, Z. Y. Han, P. Bhandari, K. B. Ghaghada, A. Badea, and C. T. Badea. 2023. ’Advanced photon counting CT imaging pipeline for cardiac phenotyping of apolipoprotein E mouse models’, PLoS One, 18: e0291733.

Allphin, A. J., R. Nadkarni, Z. Y. Han, D. P. Clark, K. B. Ghaghada, A. Badea, and C. T. Badea. 2025. ’Assessing the cardioprotective effects of exercise in APOE mouse models using deep learning and photon-counting micro-CT’, PLoS One, 20: e0320892.

Anderson, Robert J., James J. Cook, Natalie Delpratt, John C. Nouls, Bin Gu, James O. McNamara, Brian B. Avants, G. Allan Johnson, and Alexandra Badea. 2019. ’Small animal multivariate brain analysis (SAMBA)–a high throughput pipeline with a validation framework’, Neuroinformatics, 17: 451–72.

Avants, Brian B., Nicholas J. Tustison, Gang Song, Philip A. Cook, Arno Klein, and James C. Gee. 2011. ’A reproducible evaluation of ANTs similarity metric performance in brain image registration’, Neuroimage, 54: 2033–44.

Badea, A., W. Wu, J. Shuff, M. Wang, R. J. Anderson, Y. Qi, G. A. Johnson, J. G. Wilson, S. Koudoro, E. Garyfallidis, C. A. Colton, and D. B. Dunson. 2019. ’Identifying Vulnerable Brain Networks in Mouse Models of Genetic Risk Factors for Late Onset Alzheimer’s Disease’, Front Neuroinform, 13: 72.

Badea, Alexandra, Didong Li, Andrei R. Niculescu, Robert J. Anderson, Jacques A. Stout, Christina L. Williams, Carol A. Colton, Nobuyo Maeda, and David B. Dunson. 2022. ’Absolute winding number differentiates mouse spatial navigation strategies with genetic risk for Alzheimer’s disease’, Frontiers in neuroscience, 16: 848654.

Badea, Alexandra, Ali Mahzarnia, Divya Reddy, Zijian Dong, Robert J. Anderson, Hae Sol Moon, Jacques A. Stout, Janai Williams, Lydiane Hirschler, and Emmanuel L. Barbier. 2025. ’Neuroimaging biomarkers of neuroprotection: Impact of voluntary versus enforced exercise in Alzheimer’s disease models’, Magnetic resonance imaging, 121: 110406.

Bauer, Ulrich. 2021. ’Ripser: efficient computation of Vietoris–Rips persistence barcodes’, Journal of Applied and Computational Topology, 5: 391–423.

Bell, Robert D., Ethan A. Winkler, Itender Singh, Abhay P. Sagare, Rashid Deane, Zhenhua Wu, David M. Holtzman, Christer Betsholtz, Annika Armulik, and Jan Sallstrom. 2012. ’Apolipoprotein E controls cerebrovascular integrity via cyclophilin A’, Nature, 485: 512–16.

Belloy, M. E., V. Napolioni, and M. D. Greicius. 2019. ’A Quarter Century of APOE and Alzheimer’s Disease: Progress to Date and the Path Forward’, Neuron, 101: 820–38.

Blumenfeld, Jessica, Oscar Yip, Min Joo Kim, and Yadong Huang. 2024. ’Cell type-specific roles of APOE4 in Alzheimer disease’, Nature Reviews Neuroscience, 25: 91–110.

Cechetto, D. F. 2014. ’Cortical control of the autonomic nervous system’, Exp Physiol, 99: 326–31.

Crestani, Carlos C., Fernando H. F. Alves, Felipe V. Gomes, Leonardo Resstel, Fernando Correa, and James P. Herman. 2013. ’Mechanisms in the bed nucleus of the stria terminalis involved in control of autonomic and neuroendocrine functions: a review’, Current neuropharmacology, 11: 141–59.

Critchley, Hugo D., Christopher J. Mathias, Oliver Josephs, John O’Doherty, Sergio Zanini, Bonnie-Kate Dewar, Lisa Cipolotti, Tim Shallice, and Raymond J. Dolan. 2003. ’Human cingulate cortex and autonomic control: converging neuroimaging and clinical evidence’, Brain, 126: 2139–52.

Hirschler, Lydiane, Clément S. Debacker, Jérôme Voiron, Sascha Köhler, Jan M. Warnking, and Emmanuel L. Barbier. 2018. ’Interpulse phase corrections for unbalanced pseudo-continuous arterial spin labeling at high magnetic field’, Magnetic Resonance in Medicine, 79: 1314–24.

Lein, Ed S., Michael J. Hawrylycz, Nancy Ao, Mikael Ayres, Amy Bensinger, Amy Bernard, Andrew F. Boe, Mark S. Boguski, Kevin S. Brockway, and Emi J. Byrnes. 2007. ’Genome-wide atlas of gene expression in the adult mouse brain’, Nature, 445: 168–76.

Lin, Yongkang, Yanhui Ding, Shulei Chang, Xinting Ge, Xiaodan Sui, and Yanyun Jiang. 2024. ’RS2-Net: An end-to-end deep learning framework for rodent skull stripping in multi-center brain MRI’, Neuroimage, 298: 120769.

Montagne, A., D. A. Nation, A. P. Sagare, G. Barisano, M. D. Sweeney, A. Chakhoyan, M. Pachicano, E. Joe, A. R. Nelson, L. M. D’Orazio, D. P. Buennagel, M. G. Harrington, T. L. S. Benzinger, A. M. Fagan, J. M. Ringman, L. S. Schneider, J. C. Morris, E. M. Reiman, R. J. Caselli, H. C. Chui, J. Tcw, Y. Chen, J. Pa, P. S. Conti, M. Law, A. W. Toga, and B. V. Zlokovic. 2020. ’APOE4 leads to blood-brain barrier dysfunction predicting cognitive decline’, Nature, 581: 71–76.

Moore, Elizabeth E., Panpan Zhang, Omair A. Khan, Dandan Liu, Deepak K. Gupta, Kimberly R. Pechman, Katherine A. Gifford, Bennett A. Landman, Timothy J. Hohman, and Angela L. Jefferson. 2026. ’Lower cardiac output is a risk factor for faster cerebral atrophy over a 11-year follow-up period in APOE-ε4 carriers’, Alzheimer’s & Dementia, 22: e71598.

Nadkarni, Rohan, Alex J. Allphin, Darin P. Clark, Yi Qi, Zay Yar Han, Ketan B. Ghaghada, Alexandra Badea, and Cristian T. Badea. 2025. ’Exercise mitigates high-fat diet-induced cardiac dysfunction via APOE genotype-and immune-dependent mechanisms: A photon-counting CT study in adult mice’, PLoS One, 20: e0339293.

Safieh, M., A. D. Korczyn, and D. M. Michaelson. 2019. ’ApoE4: an emerging therapeutic target for Alzheimer’s disease’, BMC Med, 17: 64.

Saltarelli, Gaspare, Giovanni Di Cerbo, Antonio Innocenzi, Claudia De Felici, Alessandra Splendiani, and Ernesto Di Cesare. 2025. ’Quantitative MRI in neuroimaging: a review of techniques, biomarkers, and emerging clinical applications’, Brain Sciences, 15: 1088.

Schneider, Sebastian C., Stephan Kaczmarz, Jens Göttler, Jan Kufer, Benedikt Zott, Josef Priller, Michael Kallmayer, Claus Zimmer, Christian Sorg, and Christine Preibisch. 2023. ’Stronger influence of systemic than local hemodynamic-vascular factors on resting-state BOLD functional connectivity’, Neuroimage, 281: 120380.

Shu, Juan, Rong Zheng, Julio Chirinos, Carlos Copana, Bingxuan Li, Zirui Fan, Xiaochen Yang, Yilin Yang, Xiyao Wang, and Yujue Li. 2025. ’Inferring multi-organ genetic connections using imaging and clinical data through Mendelian randomization’, Nature biomedical engineering: 1–16.

Songdechakraiwut, Tananun, and Moo K. Chung. 2020. "Dynamic Topological Data Analysis for Functional Brain Signals." In. Stolz, B. J., H. A. Harrington, and M. A. Porter. 2017. ’Persistent homology of time-dependent functional networks constructed from coupled time series’, Chaos, 27: 047410.

Valenza, Gaetano, Zoran Matić, and Vincenzo Catrambone. 2025. ’The brain–heart axis: integrative cooperation of neural, mechanical and biochemical pathways’, Nature Reviews Cardiology, 22: 537–50.

Vorhees, Charles V., and Michael T. Williams. 2006. ’Morris water maze: procedures for assessing spatial and related forms of learning and memory’, Nature protocols, 1: 848–58.

Wang, Quanxin, Song-Lin Ding, Yang Li, Josh Royall, David Feng, Phil Lesnar, Nile Graddis, Maitham Naeemi, Benjamin Facer, and Anh Ho. 2020. ’The Allen mouse brain common coordinate framework: a 3D reference atlas’, Cell, 181: 936–53.

Winter, S., A. Mahzarnia, R. J. Anderson, Z. Y. Han, J. Tremblay, J. A. Stout, H. S. Moon, D. Marcellino, D. B. Dunson, and A. Badea. 2024. ’Brain network fingerprints of Alzheimer’s disease risk factors in mouse models with humanized APOE alleles’, Magn Reson Imaging, 114: 110251.

Yamazaki, Y., N. Zhao, T. R. Caulfield, C. C. Liu, and G. Bu. 2019. ’Apolipoprotein E and Alzheimer disease: pathobiology and targeting strategies’, Nat Rev Neurol, 15: 501–18.

Zalesky, Andrew, Ye Ella Tian, Luigi Ferrucci, Wenjia Bai, Michael S. Rafii, Paul Aisen, Jian Zeng, Aleix Boquet-Pujadas, Filippos Anagnostakis, Michael R. Duggan, Cassandra M. Joynes, Arthur W. Toga, Zhijian Yang, Keenan A. Walker, Christos Davatzikos, Junhao Wen, and Multi Consortium The. 2026. ’Brain–heart–eye axis revealed by multi-organ imaging genetics and proteomics’, Nature biomedical engineering, 10: 1171–93.

Zerbi, V., M. Wiesmann, T. L. Emmerzaal, D. Jansen, M. Van Beek, M. P. Mutsaers, C. F. Beckmann, A. Heerschap, and A. J. Kiliaan. 2014. ’Resting-state functional connectivity changes in aging apoE4 and apoE-KO mice’, J Neurosci, 34: 13963–75.

Zhao, B., T. Li, Z. Fan, Y. Yang, J. Shu, X. Yang, X. Wang, T. Luo, J. Tang, D. Xiong, Z. Wu, B. Li, J. Chen, Y. Shan, C. Tomlinson, Z. Zhu, Y. Li, J. L. Stein, and H. Zhu. 2023. ’Heart-brain connections: Phenotypic and genetic insights from magnetic resonance images’, Science, 380: abn6598.

