## SupplementaryMaterial for "APOE isoforms shape brain network organization and its coupling with cardiac phenotypes in mice"

### Supplementary Methods S2 | Functional network analyses

#### Methods

#### Cohort and imaging-derived networks

The functional-connectivity analysis comprised 292 mice: 109 APOE2, 69 APOE3, 83 APOE4 and 31 Apoe knockout (KO). The primary humanized APOE contrasts and persistent-homology analysis included 261 mice. After applying the common-availability mask and restricting to gray-matter atlas nodes, 244 regions (29,646 undirected edges) were retained. Subject-level connectivity matrices were taken from the saved windowed functional-connectivity archive; upper-triangular entries were used for the multivariate connectome analysis. Mice with flagged or pending metadata, missing protocol assignment, or incomplete required covariates were excluded according to the analysis scripts.

#### Network measures, regional strength and genotype inference

Nine global functional-network outcomes were tested, including positive and negative mean strength, global efficiency, clustering, modularity, and positive and negative $\beta_0$ and $\beta_1$ persistence AUCs. The primary network-organization measures displayed in panel A were positive strength, clustering and modularity. Outcomes were rank transformed and fitted to covariate-adjusted regression models. Regression effects are reported as standardized coefficients on the ranked outcome scale, with 95% bootstrap confidence intervals. Permutation p values used 5,000 permutations and effect intervals used 2,000 bootstrap replicates in the final genotype/risk-factor/KO analyses. The implementation used the model's covariate-adjustment and permutation procedures; nominal and FDR-adjusted p values must be interpreted in their respective prespecified test families rather than compared across differently scoped analyses.

Regional positive and negative node strengths were recomputed from the windowed functional-connectivity matrices on the 244-node common-coverage gray-matter network (at least one EPI-voxel equivalent in at least 95% of mice passing registration quality control). The six original zero-variance regions were among 16 gray-matter regions excluded by this prespecified coverage criterion. For each node, sign-specific strength was the sum of positive edge weights or the absolute magnitudes of negative edge weights, divided by 243. The 261-mouse, corrected-body-mass regional analysis used rank-transformed strengths in covariate-adjusted ordinary least squares models, without exercise. Two-sided t-test p values were adjusted with Benjamini–Hochberg (BH) FDR across 244 nodes separately for each of three APOE genotype contrasts and each connection sign (six regional testing families). Findings with $q < 0.05$ were considered significant; $0.05 \le q < 0.10$ was considered suggestive. The regional analysis was rerun on this 244-node network, not obtained by filtering the earlier 260-node coefficients.

#### Cross-validated connectome embedding and network-measure PCA

Per-edge genotype/group effects were estimated in covariate-adjusted linear models using APOE2 as the reference. To reduce noise-induced positive bias in group centroid distances, groups were randomly divided into two halves and cross-validated squared distances were calculated as the dot product of independently estimated group-effect differences, divided by the number of edges. Distances were averaged over 200 splits. Nonpositive squared-distance estimates were treated as having no detectable separation for visualization. Classical multidimensional scaling (MDS) was applied to the nonnegative distance matrix. Uncertainty around map coordinates was assessed using 500 within-group bootstrap resamples, with bootstrap maps aligned to the original embedding by Procrustes transformation. The global permutation test summed distances over the three humanized

APOE pairs; genotype labels were permuted among humanized mice (KO fixed), with 999 permutations.

For the network-measure PCA, covariate contributions were removed from each of nine ranked network outcomes while retaining genotype-group effects. The residualized outcome matrix was standardized and subjected to singular-value-decomposition PCA. Individual mice and group centers were plotted in the first two principal-component dimensions. Ellipses denote uncertainty in bootstrapped group centroids, not prediction regions for individual mice.

#### Persistent-homology curves and bootstrap intervals

For each of the 261 humanized APOE mice, positive and negative connectivity ranks were used to form separate sign-specific filtrations. Persistent-homology diagrams were calculated in dimensions 0 (connected components) and 1 (loops) using Ripser, then converted to Betti curves over edge density. The density grid was dense through 0.10 (step 0.0005), with additional coarser sampling extending to density 1. For each diagram, AUC was calculated as the sum of feature lifetimes truncated at density 1. Genotype-adjusted Betti curves were fitted using the same design covariates as the connectome analysis; 2,000 resamples within genotype produced pointwise 95% percentile bootstrap intervals. Saved persistence diagrams were verified against matrix SHA-256 hashes before reusing them.

The panels show only informative density ranges (0–0.10 for positive and negative $\beta_0$, 0–0.50 for positive $\beta_1$ and 0–0.80 for negative $\beta_1$), whereas annotated significance tests summarize full-filtration AUC. Pointwise bands do not represent simultaneous intervals or genotype hypothesis tests. Nonpositive adjusted $\beta_0$ values were masked only for plotting on logarithmic axes; the underlying saved numerical values were unchanged.

Supplementary Table S2A. Regional APOE contrasts and multiplicity families

| Inference family | BH q < 0.05 discoveries | Interpretation |
|---|---|---|
| APOE4 vs APOE2: positive regional strength; 244 tests | 15 | All significant estimates are negative |
| Six separate genotype-contrast × sign families; 244 tests each | 17 | Within-family BH discoveries across 1,464 tested associations |
| Pooled across all three contrasts and both signs; 1,464 tests | 1 | Left cingulate cortex area 24b; $\beta = -0.653$; $p = 2.6876 \times 10^{-5}$; $q_{244} = 0.0065576$; pooled $q_{1464} = 0.03935$ |

### Supplementary Methods S3 | Cardiac–functional-network analyses

#### Methods

#### Cohort selection, cardiac CT phenotypes, and functional connectivity measures

The frozen primary cohort contained 80 historical and 28 new-2026 mice maintained on control diet with absolute CT–MRI interval ≤90 days. Regional signed strength was derived from Fisher-z-transformed 244×244 functional connectivity matrices as the mean positive or absolute negative edge weight across 243 partner regions, with diagonal connections excluded. Network persistence outcomes included negative-edge $\beta_0$ AUC. The model included cardiac exposure × APOE genotype interaction, sex, age at MRI, body mass at CT, HN, signed CT–MRI interval, and identifiable cohort, scan-year, and protocol terms. Model terms and coding were taken from the frozen specifications. Interaction significance was assessed with HC3-robust joint two-degree-of-freedom Wald tests. Primary cardiac exposures were diastolic LA volume, diastolic myocardial volume, LV ejection fraction, and cardiac output.

#### Interaction models and multiplicity control

Global multiplicity: BH over four cardiac exposures for each topology outcome ($q_4$), and across 12 persistence-topology tests ($q_{12}$), as in the frozen primary analysis. Regional multiplicity: BH over 488 signed regional-strength outcomes within each cardiac phenotype and across 1,952 tests for the original four-metric screen; for the exploratory expanded screen, BH within each 488-test phenotype family and across all 14 × 488 = 6,832 tests. HC3-robust joint 2-df APOE × cardiac Wald tests were used, and individual contrasts and LOO significance are nominal. The four original cardiac analyses were reused unchanged and ten additional cardiac exposures were harmonized to historical units before fitting; 2026 µL volumes were divided by 1,000 to obtain mL. The atlas-size sensitivity families represent post hoc restricted tests and must not be interpreted as confirmatory. The extended 14-phenotype analysis was exploratory, and within-phenotype BH results do not imply significance under the pooled 6,832-test family.

#### Cardiac phase alignment and harmonization across acquisition eras

Cardiac harmonization: Historical diastolic LA volume was defined as maximum LA volume and harmonized to the corresponding 2026 CT_LA_Max_uL measurement. All 108 cardiac records were matched (80 historical, 28 new 2026), and the four frozen exposures were numerically reproduced. Volumes recorded in µL in 2026 were divided by 1,000 for historical mL units. The phase equivalence of historical and 2026 systolic atrial measurements remains provisional, and the median diastolic RA volume differs across cohorts (0.01077 versus 0.02251 mL). Consequently, atrial outcomes other than the confirmed LA maximum should be interpreted cautiously. Myocardial mass, calculated from myocardial volume and tissue density, is distinct from animal body mass.

**Supplementary Table S3A. Exploratory 14-phenotype screen and pooled FDR**

| Cardiac phenotype | Minimum p | Within-family BH hits | Pooled BH hits | Minimum q (within 488) | Minimum q (all 6,832) |
|---|---|---|---|---|---|
| Cardiac_Output | 0.00142 | 0 | 0 | 0.3827 | 0.5577 |
| Diastolic_LA | 6.14e-05 | 1 | 0 | 0.0299 | 0.1397 |
| Diastolic_LV_Volume | 0.000658 | 0 | 0 | 0.3212 | 0.3983 |
| Diastolic_Myo | 4.4e-05 | 1 | 0 | 0.0215 | 0.1397 |
| Diastolic_RA | 0.00227 | 0 | 0 | 0.9012 | 0.5953 |
| Diastolic_RV | 0.000198 | 0 | 0 | 0.0964 | 0.3219 |
| Ejection_Fraction | 0.00207 | 0 | 0 | 0.7405 | 0.5953 |
| Heart_Rate | 0.000283 | 0 | 0 | 0.1380 | 0.3219 |
| Stroke_Volume | 0.00118 | 0 | 0 | 0.4488 | 0.5577 |
| Systolic_LA | 0.000367 | 0 | 0 | 0.1791 | 0.3388 |
| Systolic_LV_Volume | 0.0047 | 0 | 0 | 0.9995 | 0.7646 |
| Systolic_Myo | 4.62e-05 | 1 | 0 | 0.0226 | 0.1397 |
| Systolic_RA | 0.000486 | 0 | 0 | 0.1138 | 0.3388 |
| Systolic_RV | 0.000423 | 0 | 0 | 0.2066 | 0.3388 |

Each family includes 244 regions × positive/negative connectivity strength (488 joint HC3 APOE × cardiac tests). Pooled correction includes all 6,832 tests; no pooled q<0.05 results. The two additional columns show the minimum BH-adjusted q-value among all 488 tests within each cardiac phenotype and the minimum q-value for that phenotype under pooled adjustment across all 6,832 tests. These are minima, not q-values calculated from the hit counts.

#### Supplementary Figure S3. Regional interactions and leave-one-animal-out analyses (four panels)

The four-panel regional figure presents the full N=108 cohort alongside the nested historical N=80 subset for each of two original cardiac–connectivity associations, together with single-animal-deletion sensitivity.

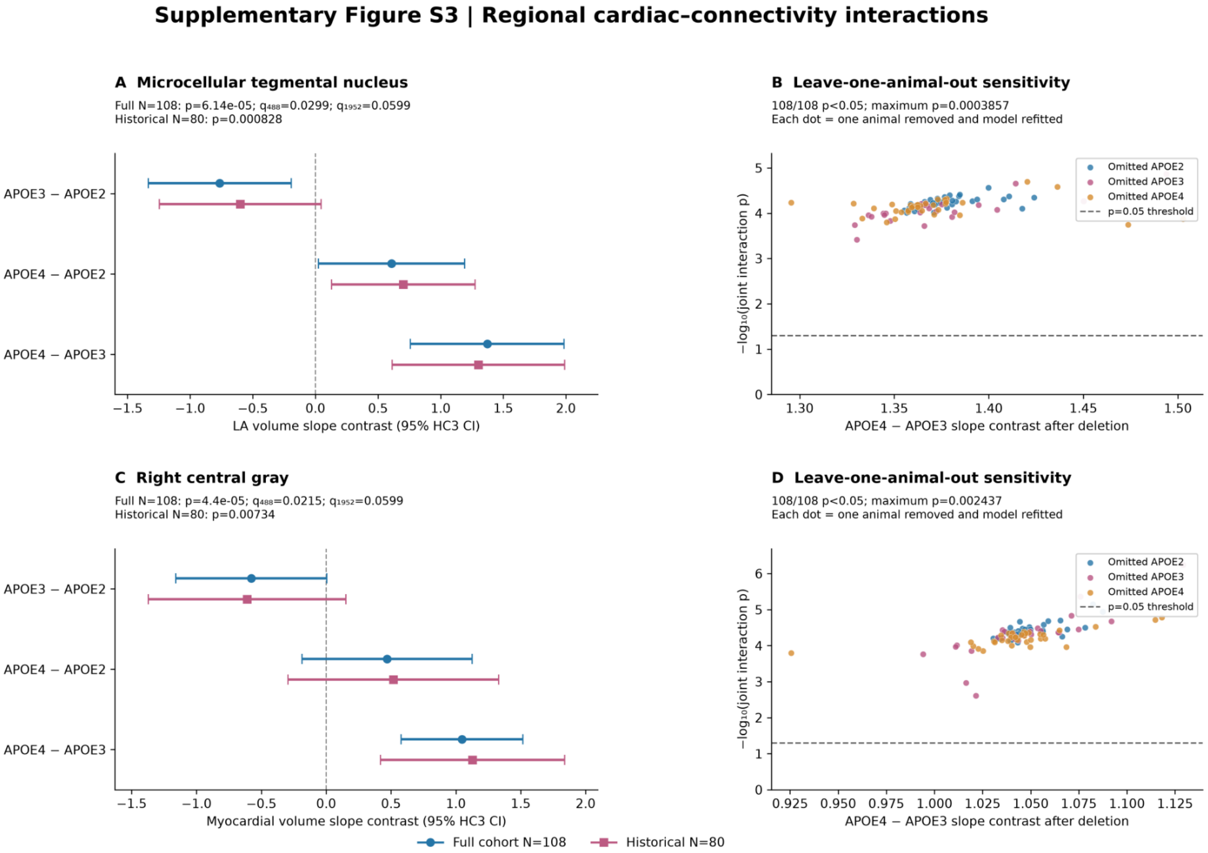


Supplementary Figure S3. Exploratory regional APOE × cardiac interactions and single-animal-deletion sensitivity. (A) Genotype slope contrasts for diastolic/maximal left atrial (LA) volume and mean positive functional connectivity strength in the microcellular tegmental nucleus, with 95% HC3-robust confidence intervals in the full cohort (N=108; blue) and nested historical subset (N=80; pink). (B) Leave-one-animal-out sensitivity for the LA interaction. Each point represents one omitted mouse; the horizontal coordinate is the APOE4−APOE3 slope contrast after deletion and the vertical coordinate is $-\log_{10}$ of the HC3-robust joint interaction p-value. Dot color identifies the genotype of the omitted mouse; the dashed reference corresponds to nominal p=0.05. All 108 refits retained p<0.05 (maximum p=0.000386). (C) Analogous genotype slope contrasts for diastolic myocardial volume and positive regional strength in right central gray. (D) Leave-one-animal-out sensitivity for the myocardial-volume interaction; all 108 refits retained nominal p<0.05 (maximum p=0.00244). The two interactions meet phenotype-specific BH FDR correction across 488 regional outcomes each ($q_{488}$=0.0299 and 0.0215, respectively), but not pooled BH correction across either the original 1,952 tests ($q_{1952}$=0.0599) or expanded 6,832 tests ($q_{6832}$=0.1397).

#### Supplementary Table S3B. Leave-one-animal-out influence

| Outcome | LOO refits | p-value range | Largest p | APOE4–APOE3 Δβ range |
|---|---|---|---|---|
| **Global negative $\beta_0$ AUC × LA** | 108/108 p<0.05 | 6.7e-07–0.0265 | 0.02648 | Retained direction; bounds not supplied |

| Outcome | LOO refits | p-value range | Largest p | APOE4–APOE3 Δβ range |
|---|---|---|---|---|
| Microcellular tegmental strength × LA | 108/108 p<0.05 | Minimum not supplied | 0.000386 | 1.295–1.503 |
| Right central gray strength × myocardial volume | 108/108 p<0.05 | 5.8e-07–0.00244 | 0.00244 | 0.926–1.129 |

All leave-one-animal-out p-values are nominal tests, not postselection FDR-adjusted significance. Retention of p<0.05 in every refit excludes dependence on any single mouse but not multiple-animal influence or confounding.

### Supplementary Methods S4 | Navigation and brain–heart–behavior analyses

#### Methods

#### Animals, design, and behavioral outcomes

Spatial navigation was evaluated in 130 mice expressing human APOE2 (n=47), APOE3 (n=44), or APOE4 (n=39). Morris water maze acquisition spanned five days, with four trials per mouse per day (2,600 acquisition trials). An acquisition winding measure was analyzed after log(1 + winding) transformation. Total swimming distance was likewise analyzed as log(1 + distance). Spatial memory was assessed with paired Day 5 and Day 8 probe measurements of target-quadrant occupancy (260 probe observations). The analysis dataset includes APOE genotype, humanized immune status (HN), sex, diet, and age at MRI.

#### Functional-connectivity predictors and regional screening

Anatomically indexed mouse-level regional functional-connectivity node-strength features were drawn from the 244-region atlas and matched to behavior using mouse identifiers. Positive and negative functional-connectivity networks were handled separately. The unbiased regional screen evaluated 244 regions × two connectivity signs × three acquisition-related endpoints (Day 1, Day 5, and learning rate), yielding 1,464 tests. Benjamini–Hochberg correction was applied across the full screen; sign/endpoint family-specific q-values are also retained in the source table. Targeted follow-up longitudinal models examined left hypothalamic PVG negative connectivity (neg_165), left cingulate area 30 negative connectivity (neg_8), and left cingulate area 25 positive connectivity (pos_149). Regional features were z-standardized. Significance within the targeted three-region models must not be conflated with significance in the 1,464-test screen.

#### Longitudinal acquisition and probe models

APOE acquisition models evaluated overall adjusted genotype effects and APOE × acquisition-day interactions, with both linear-day and categorical-day specifications. Genotype-specific acquisition slopes and Holm/BH-adjusted pairwise contrasts were reported as appropriate to their source families. The integrated regional and cardiac models used log(1 + winding), day centered at Day 3, genotype and biological covariates, and mouse-clustered uncertainty estimates. The raw acquisition trial records were checked for coverage, duplicate identifiers, and valid behavioral outcomes. Paired probe tests modeled occupancy as a function of genotype, probe day, and their interaction with mouse-clustered uncertainty; Day 5 and Day 8 pairwise genotype contrasts were Holm-corrected separately within each probe day.

#### Cardiac cohort reconciliation

Cardiac ejection fraction (EF) was linked to behavior through a curated CT-to-behavior identity ledger and cardiac identity audit. The primary conservative subset included 37 mice without the seven separately adjudicated matches; the 44-mouse analysis includes those seven and is a sensitivity analysis, not an independent replication sample. For the cardiac longitudinal models, EF was standardized within cohort; the coefficient of EF × acquisition day assessed the association between EF and day-to-day change in winding. Complete primary and sensitivity acquisition datasets contained 740 and 880 trial observations, respectively. Results were treated as observational associations, not causal effects.

#### Exploratory three-way interaction and robustness

The exploratory Panel D fit for the 37-mouse primary cohort used: log_winding ~ C(APOE) × fc_z × ef_z + C(HN) + C(Sex) + C(Diet) + age_z + day_c + trial_c, where fc_z denotes left cingulate area 25 positive FC. The interaction concerns genotype differences in the cross-sectional FC–EF association with winding, not genotype-specific acquisition learning slopes. Trial-level, mouse-day and mouse-level methods were compared. A null-restricted, mouse-level Rademacher wild-cluster bootstrap with 399 replicates assessed the joint APOE × FC × EF interaction. Within-genotype FC × EF slopes and higher-minus-lower EF contrasts at FC z=0 were extracted from the same pooled model using EF quartiles, with six-test Holm correction. This analysis is exploratory because of modest subgroup sizes and sensitivity of asymptotic inference.

##### Supplementary Table S4A. Wild-cluster bootstrap sensitivity for the three-way model

| Cohort | Unit | Mice | Obs | Asymptotic p | Bootstrap p | Valid reps |
|---|---|---|---|---|---|---|
| **PRIMARY_37** | trial | 37 | 740 | 2.27e-06 | 0.095 | 399 |
| **PRIMARY_37** | mouse_day | 37 | 185 | 5.31e-06 | 0.09 | 399 |
| **SENSITIVITY_44** | trial | 44 | 880 | 0.000924 | 0.08 | 399 |
| **SENSITIVITY_44** | mouse_day | 44 | 220 | 0.0014 | 0.0875 | 399 |

##### Supplementary Table S4B. Combined cardiac EF and cingulate connectivity acquisition model

| Cohort | Mice | Predictor | β | Cluster p | Bootstrap p | Holm p |
|---|---|---|---|---|---|---|
| **PRIMARY_37** | 37 | Cardiac EF x acquisition day | 0.0294 | 0.0314 | 0.045 | 0.09 |
| **PRIMARY_37** | 37 | Left cingulate area 25 positive FC x acquisition day | 0.0349 | 0.009 | 0.1125 | 0.1125 |
| **SENSITIVITY_44** | 44 | Cardiac EF x acquisition day | 0.0242 | 0.0247 | 0.03 | 0.06 |
| **SENSITIVITY_44** | 44 | Left cingulate area 25 positive FC x acquisition day | 0.0362 | 0.0028 | 0.0425 | 0.06 |

#### Supplementary whole-brain screening record

The complete, independently corrected regional analysis contains 1,464 results and is supplied as tables/wholebrain_longitudinal/wholebrain_longitudinal_1464.csv. The seven lowest overall q-value rows are shown here for interpretive context. The full CSV must accompany the manuscript's data supplement; the truncated display below does not substitute for all 1,464 rows.

| FC sign | Endpoint | Region | Side | β | p | Overall q |
|---|---|---|---|---|---|---|
| **neg** | Day5 | PVG_of_Hypothalamus | Left | -0.1064 | 3.42e-06 | 0.005 |
| **pos** | Day1 | Red_Nucleus_Magnocellular | Right | 0.078 | 0.000134 | 0.0982 |
| **neg** | Day5 | Cingulate_Cortex_Area_30 | Left | -0.0869 | 0.000247 | 0.0986 |
| **pos** | Day5 | Cingulate_Cortex_Area_25 | Left | 0.0996 | 0.000275 | 0.0986 |
| **neg** | Day1 | Primary_Visual_Cortex_Binocular_Area | Left | -0.0557 | 0.000337 | 0.0986 |
| **neg** | Learning_rate | PVG_of_Hypothalamus | Left | -0.0266 | 0.000505 | 0.1231 |
| **pos** | Learning_rate | Cingulate_Cortex_Area_25 | Left | 0.0296 | 0.00097 | 0.195 |

### Supplementary Methods S5 | Regional transcriptomics and molecular mapping

#### 1. Methods

##### 1.1 Imaging-derived regional APOE effect map

Resting-state functional connectivity was quantified across 244 grey-matter regions of the CHASS-based mouse atlas. For each region, the APOE4-versus-APOE2 effect on positive nodal strength was estimated with the imaging covariate model established for the connectome analyses. Regional imaging-effect maps were linked to Allen Mouse Brain Atlas in situ hybridization expression profiles for 4,082 assayed genes. An anatomically curated (hand) mapping and an independently generated registration-based mapping to the Allen Common Coordinate Framework were examined. The main anatomical scatterplot represents 89 bilateral-averaged structures; the manuscript-wide reference describes approximately 90 forebrain regions.

##### 1.2 Spatial-null and gene-set tests

For predefined cell-type marker sets and biological pathways, mean gene-wise Spearman correlations were computed between atlas expression across mapped regions and the APOE4−APOE2 nodal-strength effect map. Tests used spatially autocorrelated surrogate effect maps and size-matched random-gene-set nulls; the manuscript repository records 5,000 surrogates of each type. A combined test required both nominal null p-values to be below 0.05. Benjamini–Hochberg FDR correction was applied within the tested mapping/set family. A regional composite marker score plotted in Figure 5A is conceptually distinct from the mean of gene-wise correlation coefficients used for null inference.

##### 1.3 Cross-platform regional profiles

Laboratory RNA-seq comprised 178 regional samples from 35 humanized APOE mice spanning 12 sampled regions. Shared genes were mapped to the corresponding Allen reference regions and to a descriptive proteomics dataset (12 regions, two APOE4/HN animals, one of each sex). Regional scores were standardized within source and gene as appropriate for profile displays. The five shared excitatory neuronal markers illustrated are Slc17a7, Camk2a, Tbr1, Nrgn and Slc17a6. Across the 12 dissections, pairwise Spearman correlations summarized agreement among RNA-seq, Allen expression and proteomics for excitatory markers and the synapse GO set. These correlations assess regional expression topography, not genotype-effect replication.

#### 1.4 Cingulate genotype analyses and multiplicity

Individual-animal cingulate RNA-seq scores were evaluated in N = 32 mice (APOE2 n = 11; APOE3 n = 14; APOE4 n = 7) for 17 predefined marker/pathway signatures. The adjusted model included categorical APOE genotype, humanized NOS2 (HN) status, sequencing/processing order, sex and diet. The APOE omnibus test had 2 numerator and 25 denominator degrees of freedom. Pairwise contrasts were estimated with HC3 heteroskedasticity-robust covariance and pointwise 95% confidence intervals. Benjamini–Hochberg correction was performed over 17 APOE omnibus tests and separately over all 51 pairwise genotype contrasts. Figure 5E–H presents raw animal-level scores by genotype (non-HN circles, HN triangles); inferential statistics are from covariate-adjusted models.

#### 1.5 Sensitivity and interpretive constraints

Three significant contrasts were evaluated after restricting to non-HN mice (n = 29) and across all 32 leave-one-animal-out refits. The APOE4 expression cohort was partly confounded with sequencing/processing order, and only APOE3-versus-APOE2 is comparatively free of that imbalance. Model adjustment cannot rule out residual batch confounding. Imaging and RNA-seq animals were distinct, and the proteomics sample is too small for genotype comparisons or inferential validation.

#### Supplementary Figure S5A | Cingulate APOE omnibus molecular signatures

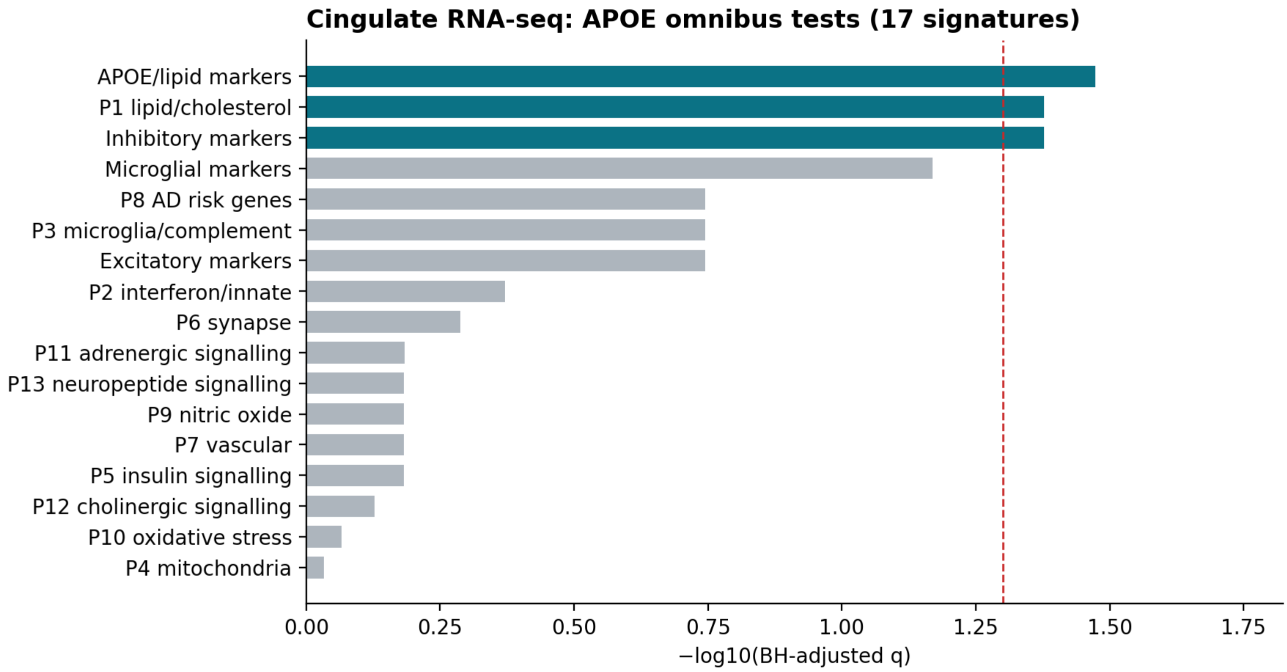


Summary of the 17 covariate-adjusted cingulate APOE omnibus tests. The dashed threshold denotes BH q = 0.05.

#### Supplementary Figure S5B | Adjusted cingulate genotype contrasts

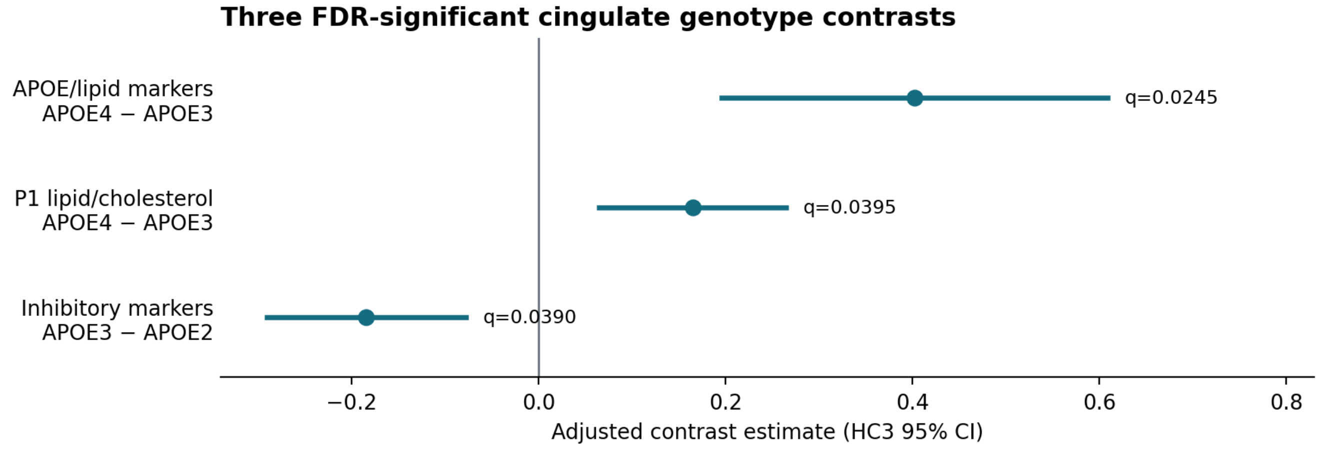


Three significant adjusted genotype contrasts with HC3 95% confidence intervals; BH correction across 51 comparisons.

#### Supplementary Table S5A. Cingulate APOE omnibus tests (17 prespecified signatures)

N = 32, df = (2,25); all models adjusted for HN, processing order, sex, diet; 17-test BH family.

| Gene set | F(2,25) | p | BH q | FDR < 0.05 |
|---|---|---|---|---|
| **Excitatory markers** | 2.89150 | 0.074195 | 0.18019 | No |
| **Inhibitory markers** | 6.35470 | 0.00587 | 0.041897 | Yes |
| **Microglial markers** | 4.90830 | 0.015919 | 0.067656 | No |
| **APOE/lipid markers** | 8.06910 | 0.0019779 | 0.033624 | Yes |
| **P1 lipid/cholesterol** | 6.00980 | 0.0073935 | 0.041897 | Yes |
| **P10 oxidative stress** | 0.21534 | 0.80774 | 0.85823 | No |
| **P2 interferon/innate** | 1.71790 | 0.19995 | 0.4249 | No |
| **P3 microglia/complement** | 3.27420 | 0.054579 | 0.18019 | No |
| **P4 mitochondria** | 0.07557 | 0.92743 | 0.92743 | No |
| **P5 insulin signalling** | 0.63367 | 0.53895 | 0.65677 | No |
| **P6 synapse** | 1.36680 | 0.27333 | 0.51628 | No |
| **P7 vascular** | 0.62993 | 0.54087 | 0.65677 | No |
| **P8 AD risk genes** | 2.94300 | 0.071157 | 0.18019 | No |
| **P9 nitric oxide** | 0.69639 | 0.50779 | 0.65677 | No |
| **P11 adrenergic signalling** | 0.99182 | 0.38503 | 0.65455 | No |
| **P12 cholinergic signalling** | 0.42625 | 0.65761 | 0.74529 | No |
| **P13 neuropeptide signalling** | 0.79623 | 0.46213 | 0.65677 | No |

#### Supplementary Table S5B. Three FDR-significant adjusted cingulate genotype contrasts (BH correction over all 51 contrasts).

| Signature | Contrast | β | 95% HC3 CI | BH q (51) |
|---|---|---|---|---|
| APOE/lipid markers | APOE4 – APOE3 | 0.40231 | 0.19577 to 0.60886 | 0.024525 |
| P1 lipid/cholesterol | APOE4 – APOE3 | 0.16492 | 0.06473 to 0.26511 | 0.039487 |
| Inhibitory markers | APOE3 – APOE2 | −0.18436 | −0.29111 to −0.07762 | 0.038996 |

**Supplementary Table S5C. Non-HN sensitivity of significant cingulate contrasts**

Nominal sensitivity-model p values are not newly family-wise corrected. Covariates: all mice = HN, processing order, sex, diet; non-HN = processing order, sex, diet.

| Signature | Contrast | Analysis | N | Estimate | SE | 95% CI | p nominal |
|---|---|---|---|---|---|---|---|
| **APOE/lipid** | E4 − E3 | All, HN-adjusted | 32 | 0.40231 | 0.10029 | [0.19577, 0.60886] | 0.00048088 |
| **APOE/lipid** | E4 − E3 | Non-HN | 29 | 0.41193 | 0.13221 | [0.13843, 0.68543] | 0.0048632 |
| **P1 lipid/cholesterol** | E4 − E3 | All, HN-adjusted | 32 | 0.16492 | 0.04865 | [0.06473, 0.26511] | 0.0023228 |
| **P1 lipid/cholesterol** | E4 − E3 | Non-HN | 29 | 0.16281 | 0.07758 | [0.00233, 0.32330] | 0.047029 |
| **Inhibitory** | E3 − E2 | All, HN-adjusted | 32 | -0.18436 | 0.05183 | [-0.29111, -0.07762] | 0.0015293 |
| **Inhibitory** | E3 − E2 | Non-HN | 29 | -0.18421 | 0.05271 | [-0.29324, -0.07518] | 0.0019518 |

**Supplementary Table S5D. Leave-one-animal-out estimate stability**

| Signature | Contrast | Full β | Fits | Fails | Min β | Max β | Sign flips | Largest \|Δβ\| |
|---|---|---|---|---|---|---|---|---|
| **APOE/lipid** | E4 − E3 | 0.40231 | 32 | 0 | 0.36500 | 0.43854 | 0 | 0.03731 |
| **P1 lipid/cholesterol** | E4 − E3 | 0.16492 | 32 | 0 | 0.14019 | 0.19163 | 0 | 0.02671 |
| **Inhibitory** | E3 − E2 | -0.18436 | 32 | 0 | -0.20752 | -0.16529 | 0 | 0.02315 |

### Supplementary Methods S6 | Regional MRI–cardiac interactions

#### Cohorts, models, and multiplicity

The regional volume cohort included 92 mice (APOE2 36, APOE3 30, APOE4 26); quantitative intrinsic T1 and CBF cohorts each included 64 mice (APOE2 35, APOE3 21, APOE4 8). Region-by-cardiac interaction models used age, sex, exercise assignment, and humanized NOS2 status, adding diet when variable in complete cases. CT-associated body mass was not included in the primary models. Standardized brain outcomes were modeled against standardized cardiac predictors interacted with APOE genotype; two-degree-of-freedom omnibus APOE × cardiac interactions used HC3-robust Wald inference. Modality-specific planned BH families comprised 4,326 volume tests and 4,312 each for intrinsic T1 and CBF. Regional APOE main effects were estimated in separate models and are reported independently of the cardiac interaction tests.

**Supplementary Table S6A. Significant regional volume–cardiac interactions**

| CHASS region | Cardiac phenotype | BH q |
|---|---|---|
| **Left Cingulate Cortex Area 24b** | Diastolic left ventricular volume | 0.02748 |
| **Right Dorsolateral Orbital Cortex** | Cardiac output | 0.00223 |
| **Right Dorsolateral Orbital Cortex** | Diastolic left ventricular volume | 0.02995 |
| **Right Dorsolateral Orbital Cortex** | Diastolic myocardial volume | 0.00223 |

| | | |
|---|---|---|
| **Right Dorsolateral Orbital Cortex** | Diastolic right ventricular volume | 0.003834 |
| **Right Dorsolateral Orbital Cortex** | Stroke volume | 0.008283 |
| **Right Dorsolateral Orbital Cortex** | Systolic myocardial volume | 0.00223 |
| **Right Dorsolateral Orbital Cortex** | Systolic right atrial volume | 0.03362 |
| **Right Dorsolateral Orbital Cortex** | Systolic right ventricular volume | 0.03362 |
| **Right Lateral Orbital Cortex** | Cardiac output | 0.005584 |
| **Right Lateral Orbital Cortex** | Diastolic right ventricular volume | 0.00223 |
| **Right Lateral Orbital Cortex** | Systolic left atrial volume | 0.03362 |

BH q values are adjusted across the full planned volume test family. For exact machine-readable precision consult TableS6_Volume_All_12_Significant_Associations.csv.

**Supplementary Table S6B. Exact T1–CBF region–cardiac overlaps**

| CHASS region | Cardiac phenotype | T1 q | CBF q |
|---|---|---|---|
| **Left Amygdalopiriform Transition Area** | Diastolic right atrial volume | 0.02618 | 0.04623 |
| **Left Bed Nucleus of the Stria Terminalis** | Systolic right ventricular volume | 0.01003 | 0.00312 |
| **Right pedunculotegmental / medial paralemniscal / supratrigeminal composite parcel** | Systolic right ventricular volume | 0.008512 | 0.0001481 |

No exact pair is also FDR significant for volume. Note that absence of a volume discovery is not an equivalence test.

**Supplementary Table S6C. Genotype-specific slopes for exact T1–CBF overlaps**

| CHASS region | MRI | APOE2 β [95% CI] | APOE3 β [95% CI] | APOE4 β [95% CI] | Interaction q |
|---|---|---|---|---|---|
| **Amygdalopiriform transition (L)** | T1 | 0.021 [-0.216, 0.259] | 0.837 [-0.297, 1.971] | 0.580 [0.246, 0.913] | 0.02618 |
| **Amygdalopiriform transition (L)** | CBF | -0.025 [-0.275, 0.224] | 0.427 [-0.441, 1.295] | 0.815 [0.396, 1.234] | 0.04623 |
| **BNST (L)** | T1 | 0.190 [-0.032, 0.413] | 0.024 [-0.267, 0.314] | 1.249 [0.698, 1.799] | 0.01003 |
| **BNST (L)** | CBF | 0.041 [-0.316, 0.399] | -0.305 [-0.945, 0.334] | 0.911 [0.557, 1.266] | 0.00312 |
| **Right pedunculotegmental / medial paralemniscal / supratrigeminal parcel** | T1 | 0.372 [0.061, 0.684] | 0.231 [-0.036, 0.498] | 1.302 [0.849, 1.755] | 0.008512 |

| | | | | | |
|---|---|---|---|---|---|
| **Right pedunculotegmental / medial paralemniscal / supratrigeminal parcel** | CBF | -0.167 [-0.483, 0.149] | -0.244 [-0.451, -0.037] | 0.757 [0.449, 1.065] | 0.0001481 |

These six rows provide the APOE2/3/4 conditional slope estimates and HC3 pointwise confidence intervals; omnibus q values test APOE × cardiac interaction, not individual slopes.

**Supplementary Table S6D. Coincidence of regional main effects and interactions**

| Modality | CHASS region | Cardiac phenotype | Interaction q | APOE main q |
|---|---|---|---|---|
| **T1** | Right Cingulate Cortex Area 29a | Cardiac output | 0.02618 | 0.000931 |
| **T1** | Left Periaquaductal Grey | Systolic right ventricular volume | 0.01801 | 0.02214 |
| **T1** | Left Reticular Nucleus of Thalamus | Systolic right ventricular volume | 4.272e-06 | 0.01246 |
| **T1** | Left Pyramidal Tract | Systolic right ventricular volume | 0.004042 | 0.006931 |
| **T1** | Left Postsubiculum | Systolic right ventricular volume | 5.71e-07 | 0.0279 |
| **T1** | Right Reticular Nucleus of Thalamus | Systolic right ventricular volume | 0.02618 | 0.03374 |
| **T1** | Right Optic Tracts | Systolic right ventricular volume | 1.434e-06 | 0.03175 |
| **T1** | Left Periaquaductal Grey | Systolic right atrial volume | 0.0219 | 0.02214 |
| **T1** | Right Motor Root of Trigeminal Nerve | Systolic right atrial volume | 0.02408 | 0.03576 |
| **CBF** | Left Fimbria | Diastolic right ventricular volume | 0.0205 | 0.04838 |

Anatomically overlapping findings are not proof of a causal neurocardiac pathway.

### Supplementary Figure S6. Expanded APOE × cardiac interaction heatmaps

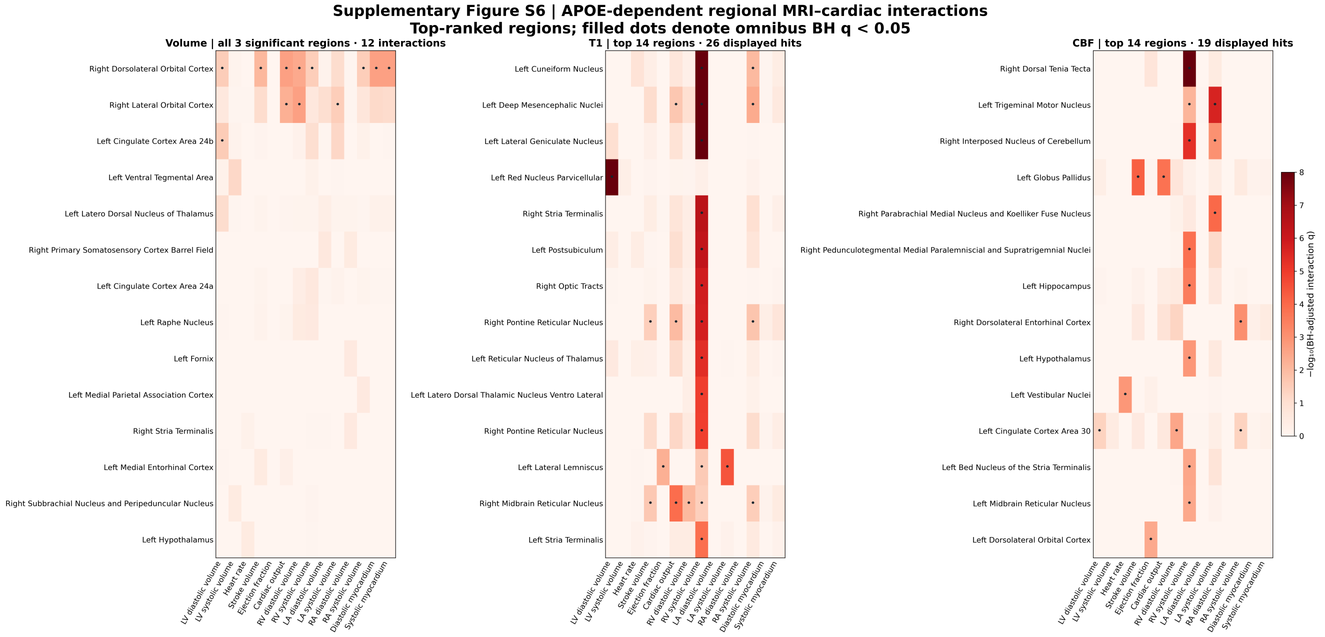


**Supplementary Figure S6. Expanded region-by-cardiac association heatmaps for regional volume, intrinsic T1 and CBF, using final HN-harmonized modality-wise FDR-corrected APOE-by-cardiac interaction tests. The full planned-test family is preserved in the accompanying CSV.**

### Supplementary Figure S7. Additional genotype-specific slopes

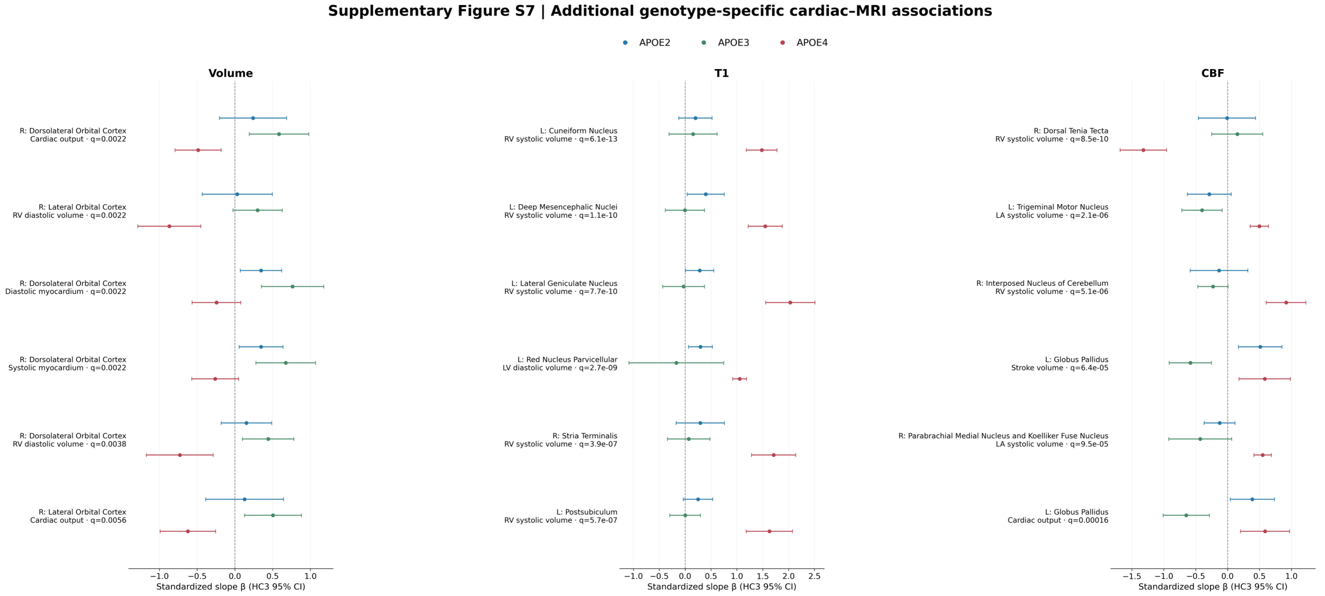


**Supplementary Figure S7. Additional conditional APOE2, APOE3, and APOE4 slopes for selected regional MRI–cardiac combinations, with HC3 95% confidence intervals. Plotted slopes must not be confused with omnibus interaction significance.**
